# Senescence-associated stromal remodeling in bone marrow aging and hematologic and bone-metastatic diseases

**DOI:** 10.64898/2026.11.15.766758

**Authors:** Julia T. Wang, Alla Karpova, Xiang Li, Kapur Dhami, Xiangwei Fang, Wen-hung Chou, Daniel R. Rapp, Preet Lal, Xiyi Wei, Yuting Zhao, Dandan Fan, Roman Smirnov, Reyka G. Jayasinghe, John M. Herndon, Andrew Houston, Cody Weimholt, Patrick Morse, Mark A. Fiala, Julie Fortier, Kiran Vij, John F. DiPersio, Russell K. Pachynski, Katherine N. Weilbaecher, Ryan Nunley, Milan Chheda, Ravi Vij, Daniel C. Link, Ryan C. Fields, Sheila A. Stewart, Feng Chen, Li Ding

**Affiliations:** Department of Medicine, Washington University in St. Louis, St. Louis, MO, 63110, USA; McDonnell Genome Institute, Washington University in St. Louis, St. Louis, MO, 63110, USA; Department of Surgery, Washington University in St. Louis, St. Louis, MO, 63110, USA; Siteman Cancer Center, Washington University in St. Louis, St. Louis, MO, 63110, USA; Department of Pathology and Immunology, Washington University in St. Louis, St. Louis, Missouri; Department of Cell Biology and Physiology, Washington University School of Medicine, St. Louis, MO 63110, USA; Bursky Center for Human Immunology and Immunotherapy, Washington University School of Medicine, St. Louis, Missouri; Department of Orthopaedic Surgery, Washington University in St. Louis, St. Louis, Missouri; Department of Cell Biology and Physiology, Washington University School of Medicine, St. Louis, MO, USA; Department of Genetics, Washington University in St. Louis, St. Louis, MO, 63110, USA

## Abstract

Cellular senescence is implicated as a driver of age-related tissue dysfunction and chronic inflammation. However, no single marker defines a senescent cell, and distinguishing senescence from other stressors requires both experimental perturbation and comparison across contexts. As part of the Cellular Senescence Network (SenNet), we generated a multimodal atlas integrating human bone marrow from 80 donors (54 assayed by single-cell RNA sequencing; 48 by Xenium spatial transcriptomics) with a 130 combined cases across four disease cohorts: breast and prostate cancer bone metastases (BCa/PCa; n=61), myeloproliferative neoplasms (MPN; n=21), and multiple myeloma (MM; n=48). Human marrow aging was characterized by interferon and IL6/JAK/STAT3 activation in hematopoietic progenitors, myeloid-biased output, and progressive loss of naïve CD4 T cell identity. Spatially, CDKN1A was concentrated in vascular stroma, and senescence-associated secretory phenotype (SASP) expression rose with age coordinately across stromal cell types within individual donors (mean pairwise ρ = 0.46), consistent with shared influences on stromal aging. Malignancy was associated with distinct and disease-specific senescence-associated states rather than uniform amplification of aging programs. Senescent fibroblasts spatially adjacent to tumor in breast and prostate cancer bone metastases shared an upregulated secretory program comprising 148 genes encoding secreted proteins, alongside upregulated matrix-remodeling and cellular adhesion signatures. Osteolineage cells found in metastasis sites as well as in MPN marrow also displayed distinct senescence-associated phenotypes, including enlarged, multinucleated osteoclasts and *CDKN1A*+ osteoblasts with upregulated secretory programs. However, senescence-associated secretory programs and matrix dysregulation in stromal cells was not observed in MM, indicating that these features are not generic consequences of tumor burden within the bone marrow but rather arise from specific cancer-stromal interactions. Residual hematopoietic regions showed altered myeloid and erythroid composition and transcriptional states. To complement these human datasets, we used a mouse model in which doxorubicin-induced senescence reproduced molecular and compositional hallmarks of marrow aging that were selectively reversible with senolytic treatment. These data establish normal marrow as a quantitative baseline for senescence and separate shared from context-specific senescent states.

## Introduction

The bone marrow (BM) microenvironment is essential for the maintenance of lifelong hematopoiesis, providing the molecular cues that regulate hematopoietic stem and progenitor cell (HSPC) maintenance, differentiation, and maturation^1^. These processes are coordinated by specialized, perivascular and CXCL12^+^ mesenchymal stroma-enriched niches that provide distinct signals to support hematopoietic function^2,3^. However, the cellular organization and molecular phenotypes of human BM stromal and progenitor populations remain incompletely characterized, due in part to their relative rarity, fragility, and limited recovery with conventional single-cell approaches. Recent enrichment-based single-cell and spatial profiling studies have begun to address this gap, revealing complex heterogeneity within the human marrow microenvironment^4^.

Aging is associated with declining BM cellularity and hematopoietic output, as well as profound alterations to its cell type composition^5^. These changes reflect not only cell-intrinsic alterations but also age-associated remodeling of the supporting niche, including loss of endosteal osteoprogenitors, expansion of central marrow mesenchymal stromal populations, and increased inflammatory signaling that can disrupt hematopoietic homeostasis^6^. In parallel, age-associated clonal hematopoiesis, driven by somatic mutations in genes such as *DNMT3A*, *TET2*, and *ASXL1*, contributes to inflammatory signaling and increased hematologic malignancy risk^7,8^. Cellular senescence represents a potential mechanistic link between these processes. Characterized by stable cell cycle arrest and a senescence-associated secretory phenotype (SASP), senescent cells can promote chronic inflammation and tissue remodeling through the secretion of cytokines, chemokines, and extracellular matrix-modifying factors^9,10^. Although senescence can act as a tumor-suppressive mechanism by limiting the proliferation of damaged or oncogenically stressed cells, persistent senescent populations may paradoxically promote tumor progression through SASP-mediated inflammation, tissue remodeling, and altered stromal signaling^11,12^. Experimental clearance of senescent cells has been shown to ameliorate age-associated pathology, including skeletal deterioration and hematopoietic dysfunction^13–15^. Despite these findings, the identity and spatial organization of putative senescent populations in human BM remain poorly defined, and it is unclear whether malignancy amplifies senescence-associated programs observed during normal aging or induces distinct forms of microenvironmental dysfunction.

In this study, we investigated age-associated remodeling and senescence-associated cellular states in the BM using complementary single-cell, spatial transcriptomic, and experimental approaches (Figure 1a). To investigate the contribution of senescence to age-associated marrow remodeling, we profiled 14 mice across four conditions: young, young plus chemotherapy, old, and old plus senolytic treatment. We next established a normal bone marrow (NBM) cohort of hematologically normal donors sampled by iliac crest biopsy or orthopedic surgery. Single-cell RNA sequencing was performed on 54 donors, 20 of which further underwent CD34+ HSPC enrichment, and Xenium spatial transcriptomics on 48 donors using either a 477-plex (n=29) or 5K whole-transcriptome panel (n=19; Figure 1b-c). Finally, we profiled four malignant cohorts using Xenium: breast (n=20, 1M cells) and prostate (n=41, 2.5M cells) cancer bone metastases, representing marrow-extrinsic tumor invasion, and myeloproliferative neoplasms (n=21, 2.3M cells) and multiple myeloma (n=48, 3.3M cells), representing marrow-intrinsic fibrotic and non-fibrotic malignancies, respectively. Together, these cohorts enabled identification of senescence-associated features responsive to senolytic treatment, characterization of normal human marrow aging, and comparison of physiological aging with disease-associated tissue stress and microenvironmental remodeling.

**Figure 1.**
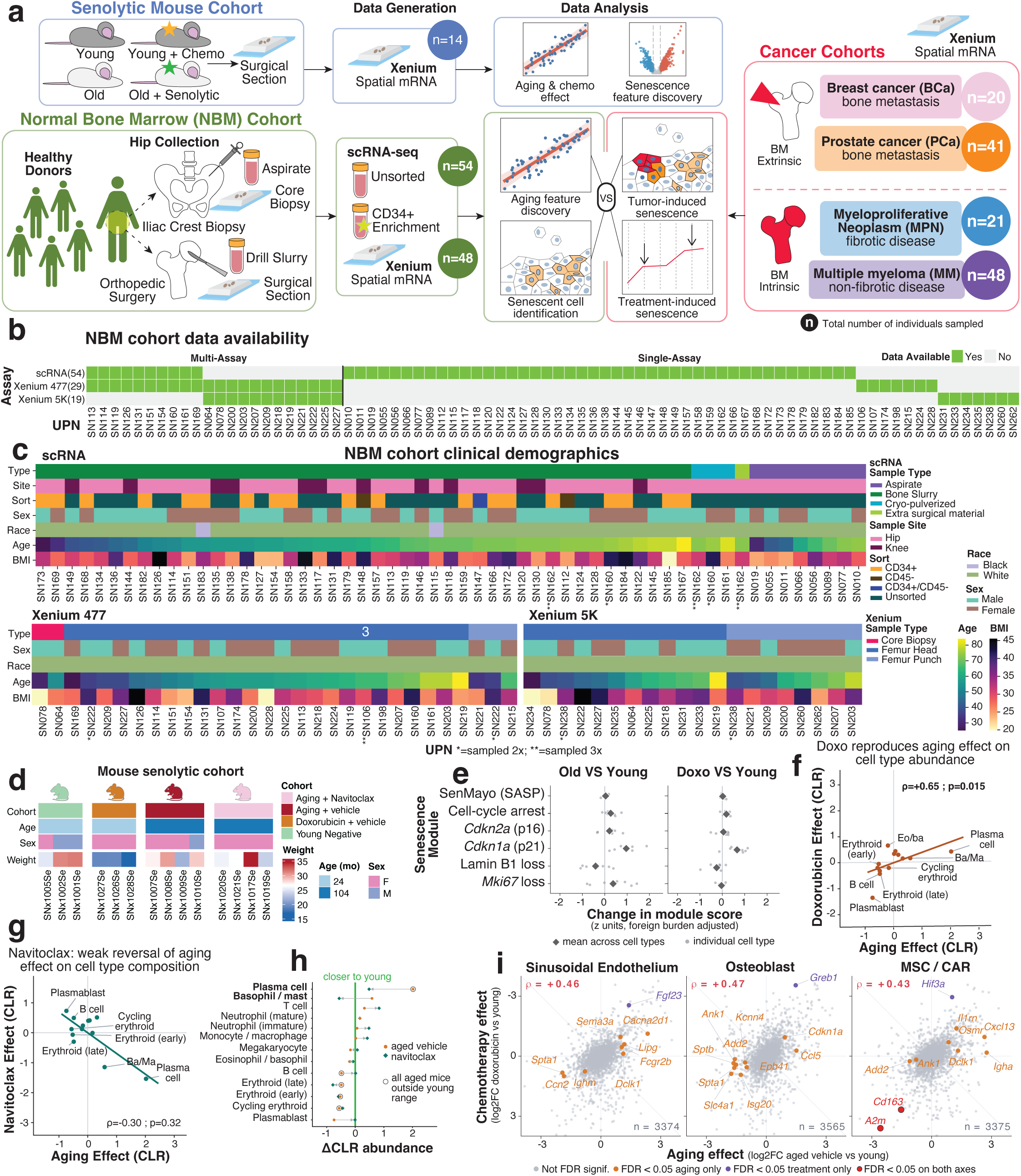
Study design and mouse senolytic experimental findings. a) Overview of study design. Number of total individuals sampled per cohort is denoted as n in colored circles. b) Data availability by assay type for NBM cohort. c) Clinical demographics of NBM cohort for scRNA-seq and Xenium datasets. d) Details of mouse experimental cohort. Navi = Navitoclax; Doxo = Doxorubicin. Aged mice and Doxorubicin-treated mice were given either vehicle control or Navitoclax; young negative controls were not given either. e) Change in senescence module across cell types in (SenMayo SASP gene set score, cell-cycle arrest gene set score, Cdkn2a or Cdkn1a expression, Lamin B1 or Mki67 loss) in old versus young and doxorubicin-treated versus young mice. f) Correlation of doxorubicin effect and aging effect (CLR) on bone marrow cell type abundance. g) Correlation of navitoclax effect and aging effect (CLR) on bone marrow cell type abundance. h) Change in CLR of cell type abundance in aged mice exposed to vehicle or Navitoclax. 0 indicates levels of young mice. i) DEGs associated with aging and/or Doxorubicin exposure in stromal cell types.

## Results

### Doxorubicin reproduces and Navitoclax reverses effects of bone marrow aging in mice

To establish a framework for investigating senescence-associated BM changes, we first asked whether experimentally induced senescence recapitulates features of physiological marrow aging and whether these changes are reversible with senolytic treatment. We addressed these questions using Xenium spatial transcriptomics in young and aged mice, where dexorubicin-induced senescence and navitoclax treatment provided complementary models of senescence induction and clearance (n = 14; Fig. 1d).

Both aging and doxorubicin induced similar senescence-associated transcriptional changes across marrow cell types (Fig. 1e), most prominently increased *Cdkn1a* (p21), with more modest increases in *Cdkn2a* (p16), cell-cycle arrest and SenMayo SASP scores, and reduced Lamin B1 and *Mki67*. The concordant magnitude and direction of these changes indicate that doxorubicin recapitulates key transcriptional features of physiological marrow aging. Cellular composition showed similar convergence. Across cell types, doxorubicin-induced changes in centered log-ratio (CLR) abundance were positively correlated with aging-associated changes (ρ = 0.65, P = 0.015; Fig. 1f). Both conditions were associated with expansion of plasma cells and basophils/mast cells and contraction of plasmablast and erythroid populations. Thus, senescence induction reproduced both the molecular and compositional features of marrow aging.

Navitoclax shifted cell composition in the opposite direction, although its effects were not significantly anticorrelated with aging across all cell types (ρ = −0.30, P = 0.32; Fig. 1g). At the individual population level, treatment shifted plasma cells and basophils/mast cells, the populations most expanded with aging, toward young levels. In contrast, erythroid and plasmablast populations remained outside the young range regardless of treatment (Fig. 1h), indicating selective rather than global reversal of age-associated compositional changes.

Finally, we examined aging and doxorubicin-associated transcriptional changes in the stromal niche, focusing on sinusoidal endothelium, osteoblasts, and MSC/CAR cells, which exhibited pronounced transcriptional alterations with aging (Fig. 1i). Although gene-level effect sizes were positively correlated between aging and doxorubicin across all three populations (ρ = 0.46, 0.47, and 0.43, respectively; n = 3,374, 3,565, and 3,375 genes), overlap in significant DEGs was limited. MSC/CAR cells showed the greatest convergence, with *A2m* and *Cd163* significantly altered under both conditions. Other aging-associated changes included increased *Cdkn1a* and *Ccl5* in osteoblasts and altered expression of inflammatory mediators, including *Il1rn*, *Osmr*, and *Cxcl13*, in MSC/CAR cells.

Together, these findings indicate that experimentally induced senescence recapitulates several molecular, compositional, and stromal features of physiological marrow aging, while senolytic treatment selectively reverses a subset of age-associated changes. Having identified senescence-associated changes in BM cell composition and inflammatory gene expression in mice, we next examined whether related changes accompany human BM aging and how they are organized across cell populations and spatial niches in an independent donor cohort.

### Increased inflammatory gene expression in progenitors, myeloid bias, and T-cell activation associated with bone marrow aging

We next characterized the cellular and transcriptional landscape of human BM aging using single-cell RNA sequencing of samples from 54 donors. Samples were obtained from iliac crest aspirates or orthopedic surgical drill slurry, and a subset was processed with CD34⁺ enrichment to improve resolution of early progenitor populations (Fig. 1a). We recovered 385,528 cells across 60 cell types and six lineages (Fig. 2a-b; Supplementary Fig. 1a-b). CD34⁺ enrichment improved resolution of early progenitors, while unsorted samples preserved native composition for abundance analyses.

**Figure 2.**
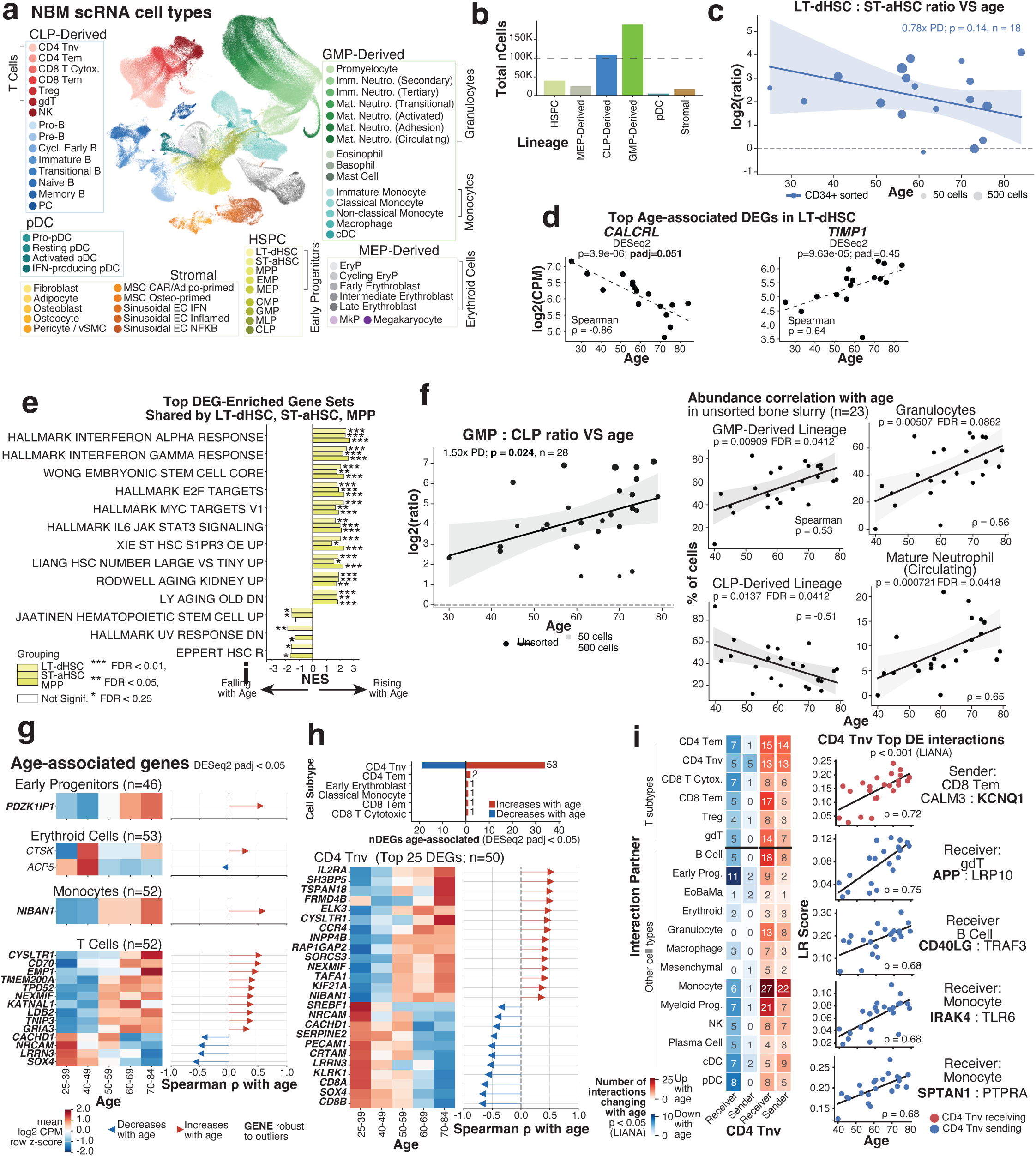
Progenitor dysregulation, myeloid bias, and T cell immunoaging in NBM scRNA-seq. a) UMAP of manually-annotated cell types captured by scRNA-seq of liquid marrow biopsies from 54 individuals. b) Total number of cells belonging to each lineage (as indicated in panel a) in dataset. c) Quasi-binomial GLM of LT-dHSC:ST-aHSC ratio with age. Dot size indicates number of LT-dHSC + ST-aHSC per donor. Abbreviations: PD=per decade. d) Correlation of log2(CPM) expression versus age of top age-associated DEGs (pseudobulk DESeq2) in LT-dHSCs. e) Top gene sets enriched by nominally-significant, age-associated DEGs (pseudobulk DESeq2), shared by at least two of LT-dHSCs, ST-aHSCs, and MPPs. NES = normalized enrichment score. f) Correlation of GMP to CLP abundance ratio and abundances of GMP-derived lineage, CLP-derived lineage, granulocyte group (denoted by bracket in panel a), and mature neutrophil (circulating) cell type with age in unsorted scRNA-seq samples. g) Age-associated genes (DESeq2 FDR<0.05; robust to outliers) in cell type groupings denoted by brackets in panel a. Heatmap on left denotes mean row z-score of log2 CPM by age bin; Spearman rho indicated on right. h) Total number of age-associated DEGs (DESeq2 FDR<0.05) in T cell types (top); mean expression and Spearman rho of age-associated DEGs in CD4 Tnv (bottom). i) Inferred age-associated ligand-receptor interaction analysis (LIANA) of CD4 Tnv. Total number of interactions changing with age, split by sender/receiver status, on left; correlation of LR score with age in top DE interactions involving CD4 Tnv on right.

HSPC abundance and composition showed trends toward age-related depletion of quiescent stem cells, although most did not reach statistical significance. Total HSPC abundance declined with age (ρ = −0.33, FDR = 0.237; Supplemental Fig. 2a), alongside a 0.78-fold decrease per decade in the ratio of long-term dormant to short-term activated hematopoietic stem cells (ST-aHSCs; p = 0.14; Fig. 2c; Supplementary Fig. 2a). Among individual subtypes, ST-aHSC expansion showed the strongest trend (OR/decade = 1.20, p = 0.075), while long-term dormant HSCs (LT-dHSC) and multipotent progenitor (MPP) abundance remained unchanged (Supplementary Fig. 2b). Few individual genes survived multiple-testing correction, although *CALCRL* declined in LT-dHSCs and *TIMP1*, a pro-inflammatory marker in disease contexts^16^, showed a strong nominal increase with age (Fig. 2d). In contrast, pathway analysis revealed a consistent inflammatory shift across LT-dHSC, ST-aHSC, and MPP, characterized by increased interferon-α/γ and IL6–JAK–STAT3 signaling and reduced quiescent HSC signatures (Fig. 2e; Supplementary Fig. 2c).

Hematopoiesis shifted toward myeloid output, with the granulocyte-monocyte progenitor (GMP): common lymphoid progenitor (CLP) ratio increasing 1.50-fold per decade (p = 0.024; Fig. 2f). Consistent with this shift, GMP-derived populations increased while CLP-derived populations declined (FDR = 0.0412 for both), driven predominantly by granulocytes, particularly mature neutrophils (ρ = 0.65, FDR = 0.0418), with concordant trends across neutrophil maturation states (Supplementary Fig. 2d).

Age-associated transcriptional remodeling was most pronounced in T cells and localized almost entirely to naïve CD4 T cells. Whereas early progenitors, erythroid cells, and monocytes showed few significant age-associated genes, naïve CD4 T cells exhibited 53 genes increasing with age; all other T-cell subtypes had at most two (Fig. 2g,h). This signature reflected progressive loss of naïve identity and acquisition of activated features, including increased *IL2RA* and *CCR4* and decreased *LRRN3* and *SOX4*^17–19^. Naïve CD4 T cells also exhibited the most extensive age-associated remodeling of cell-cell communication, with increased interactions primarily involving monocytes and myeloid progenitors, followed by B cells. Loss of inferred communication primarily involved early progenitors (Fig. 2i; Supplementary Fig. 2e).

Together, these findings suggest that within the hematopoietic department, human marrow aging is primarily characterized by inflammatory response in HSPCs, myeloid-skewed hematopoietic output, and progressive transcriptional remodeling of naïve CD4 T cells.

### Coordinated increase in stromal SASP signatures associated with bone marrow aging

To resolve the spatial organization of senescence-associated cells, we profiled BM sections using Xenium in situ transcriptomics. The 477-plex panel captured 2,221,723 cells across 23 cell types from 29 donors (Fig. 3a), annotated by marker expression following H&E-based tissue quality control and segmentation staining (Fig. 3b; Supplementary Fig. 3a). Age-associated compositional changes were modest, with trends toward increased vSMC/pericytes and decreased early B cells (Supplementary Fig. 3b).

**Figure 3.**
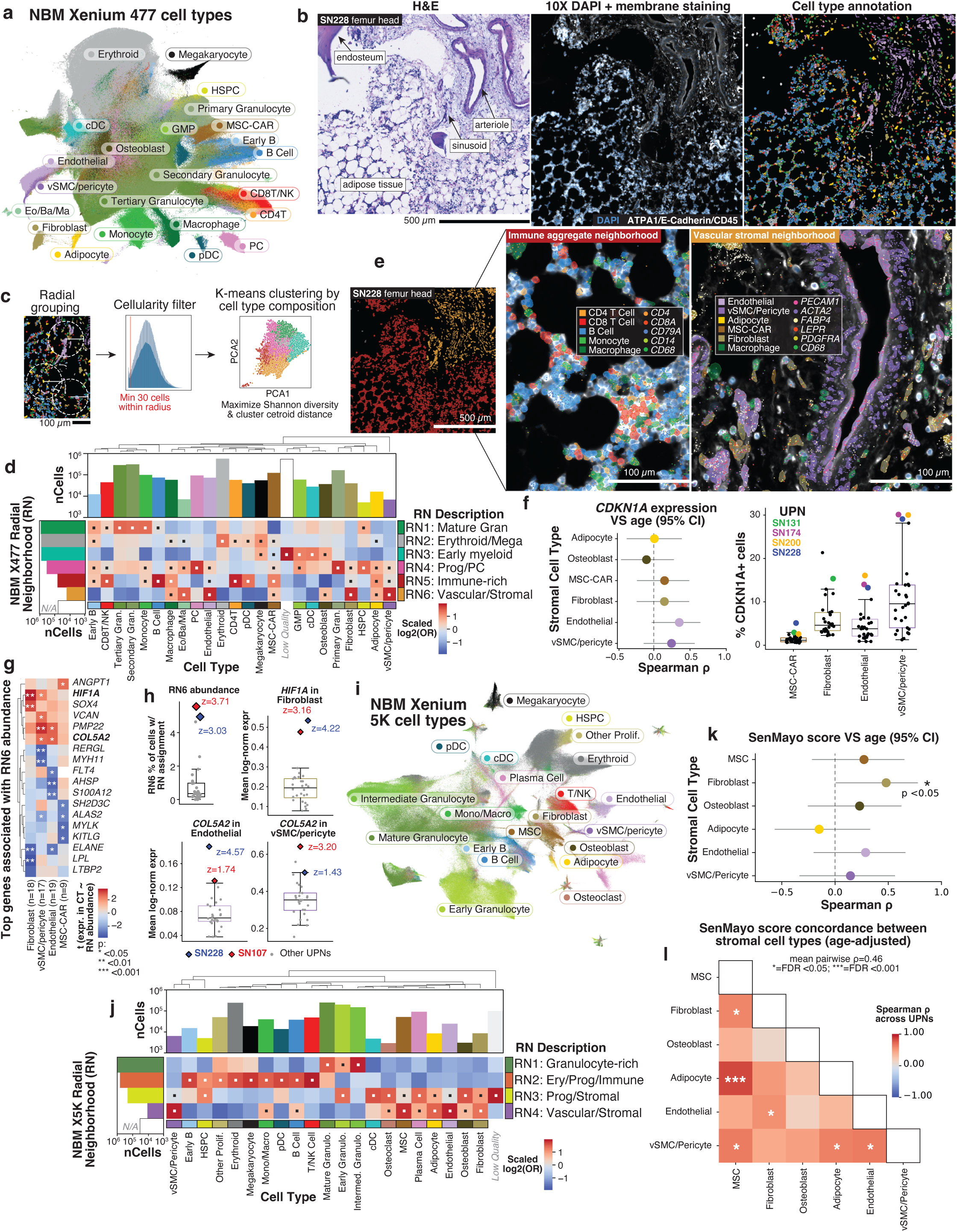
NBM spatial organization and evidence of coordinated stromal senescence with age. a) UMAP of Xenium 477 cell types (29 individuals sampled) b) Xenium spatial data processing workflow: histological H&E used for sample QC and selection; 10X segmentation staining; cell type annotation using marker gene expression (cluster-based; colors consistent with those in panel a). c) Overview of Xenium radial neighborhood detection. Following cell type annotation, clustering standardized neighbor-fraction vectors derived from all cells within 100 µm of each focal cell (minimum of 30 within-radius neighbors for inclusion) yielded 6 KMeans-defined ‘radial neighborhoods’ (RN) selected by optimizing a composite clustering metric. d) Cell-type enrichment of Xenium 477 RNs quantified using odds ratios with Fisher’s exact test followed by BH FDR correction; barplot of total cell count in dataset belonging to each RN on right and total cell count belonging to each cell type on top. e) Representative images depicting cell type breakdown within two different RNs. Colored overlays on segmented cells denote cell type; marker transcripts are visualized as colored dots. f) Mean and 95%CI of Spearman correlation *CDKN1A* expression and age in stromal cell types (left); % *CDKN1A*+ in stromal cell types (right) with UPNs high for multiple cell types highlighted. g) Top genes associated with RN6 abundance in stromal cell types. Heatmap is colored by t score of expression in cell type ∼ RN6 abundance. h) Distribution of RN6 abundance, *HIF1A* expression in fibroblasts, and *COL5A2* expression in endothelial cells and vSMC/pericytes with SN228 and SN107 UPNs highlighted. i) UMAP of Xenium 5K cell types (19 individuals sampled) j) Cell-type enrichment of Xenium 5K RNs quantified using odds ratios with Fisher’s exact test followed by BH FDR correction; barplot of total cell count in dataset belonging to each RN on right and total cell count belonging to each cell type on top. k) SenMayo score versus age (95% CI) in stromal cell types. l) Spearman correlation of SenMayo scores between stromal cell type pairs.

To characterize tissue architecture independently of cell-type annotation, we clustered cells by the composition of their 100 µm radial neighborhoods, retaining cells with at least 30 neighbors (Fig. 3c). This identified six radial neighborhoods (RNs) corresponding to granulocyte (RN1), erythroid/megakaryocyte (RN2), early myeloid (RN3), progenitor/plasma cell (RN4), immune-rich (RN5), and vascular/stromal compartments (RN6), which were recognizable as distinct structures in situ (Fig. 3d,e).

We next examined the spatial distribution of senescence markers. *CDKN1A* expression showed positive age associations in endothelium and vSMC/pericytes and a negative association in osteoblasts, although none reached significance (Fig. 3f). However, *CDKN1A* cells were substantially more abundant in vascular stroma than in other stromal populations, independent of sequencing depth (Fig. 3f; Supplementary Fig. 3c). *CDKN1A* fractions also tended to increase across multiple stromal populations within individual donors, suggesting coordinated variation in senescence burden.

Consistent with this pattern, genes associated with RN6 (vascular/stromal) neighborhood abundance showed shared changes across stromal populations, including *HIF1A* and *COL5A2* (Fig. 3g). Two donors, SN228 and SN107, exhibited particularly high RN6 abundance alongside elevated *HIF1A* in fibroblasts and *COL5A2* in endothelium and vSMC/pericytes (z = 1.4–4.6; Fig. 3h), linking vascular niche expansion with hypoxic and matrix-remodeling programs.

To validate these spatial patterns across platforms, we profiled an overlapping donor set using the Xenium Prime 5K panel, capturing 1,251,640 cells across 20 cell types from 19 donors (Fig. 3i; Supplementary Fig. 3d). Radial neighborhood analysis recovered four similarly organized neighborhoods, including a distinct vascular/stromal compartment (Fig. 3j). Cell-type proportions were highly concordant among the 12 donors profiled on both platforms (Pearson r = 0.86, Spearman ρ = 0.88), with residual differences reflecting panel-specific annotation resolution (Supplementary Fig. 3e).

SenMayo SASP scores increased with age across most stromal populations in the 5K dataset, reaching significance in fibroblasts (Fig. 3k). After adjusting for age, SASP scores remained correlated across stromal cell types within donors (mean pairwise ρ = 0.46), with significant correlations in six cell-type pairs, including MSC–fibroblast, MSC–adipocyte, and endothelial– vSMC/pericyte (Fig. 3l). These findings indicate coordinated SASP-associated expression across the stromal compartment, consistent with shared microenvironmental influences on stromal aging.

### Metastatic cancer triggers fibrosis featuring extensive senescence and unique SASP

Spatial transcriptomic analysis of breast (BCa) and prostate (PCa) cancer bone metastases identified distinct fibroblast, MSC, and senescent fibroblast states (Fig. 4a). Senescent fibroblasts exhibited cancer-associated fibroblast (CAF) matrix-remodeling programs alongside elevated *CDKN1A*, DNA damage-response genes, and known SASP markers, with limited proliferation-marker expression (Fig. 4b). Senescent fibroblasts also showed extensive upregulation of secreted genes, including 173 in breast and 183 in prostate cancer, exceeding the corresponding numbers in other fibroblast states (Fig. 4c). The 148 shared secreted genes included 20 members of the SenMayo signature, such as *CXCL12*, *KITLG*, *MMP9*, and *SERPINE1*, and were expressed more strongly in the two senescent fibroblast groups than in the other stromal reference groups, including MPN and MM fibroblasts and MSCs (Fig. 4d).

**Figure 4.**
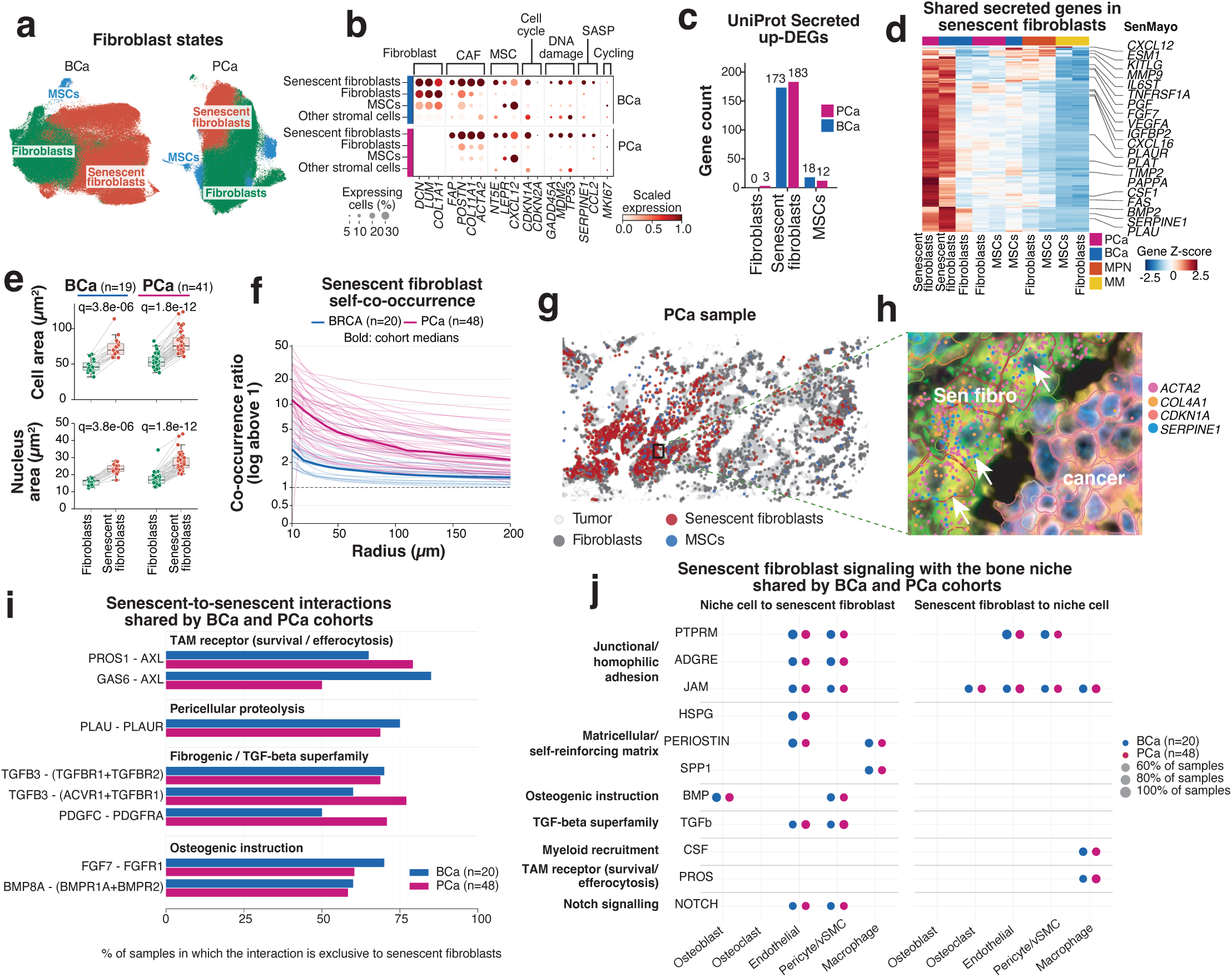
Metastatic cancer triggers fibrosis featuring extensive senescence and unique SASP. a) UMAP of fibroblasts and MSCs from BCa (left) and PCa (right) cohorts colored by their state. b) Dot plot showing expression of fibroblast and MSC marker genes, cell cycle arrest genes, DNA damage response genes and SASP gene expression in fibroblast and MSC states and other stromal cells pooled together. c) Barplot showing count of significantly up-regulated secreted DEGs (Wilcoxon rank-sum test FDR <0.05**, **log2FC >0.25, and expression in ≥5% of cells) d) Heatmap showing up-DEGs from panel c uniquely shared by senescent fibroblasts. e) Box plots comparing cell area and nucleus area between senescent and non-senescent fibroblasts in BCa and PCa cohorts. f) Xenium-based senescent fibroblast self-co-occurrence analysis showing co-localization with themselves at short distances. g) Example clustered distribution of fibroblast and MSC states in a PCa sample. Red dots show senescent fibroblasts, blue-MSCs, dark grey - regular fibroblasts, light gray - cancer cells. h) Zoom-in view of clustered senescent fibroblasts with marker gene expression. White arrows point to several senescent fibroblasts. Red outline marks all senescent fibroblasts, yellow outline marks cancer cells i) Senescent-to-senescent (autocrine) axes exclusive to the senescent state in ≥50% of samples in both cohorts, grouped by functional theme. Both cohorts independently engage TAM-receptor survival signalling (GAS6/PROS1–AXL), pericellular proteolysis (PLAU–PLAUR), TGF-β superfamily fibrogenic signalling, and osteogenic instruction (FGF7–FGFR1, BMP8A– BMPR1A/BMPR2). j) Signalling between senescent fibroblasts and the marrow niche, shown for the five cell types annotated in both cohorts, split by direction: niche cell to senescent fibroblast (left) and senescent fibroblast to niche cell (right). Twenty-four pathway-family × partner × direction axes reached ≥60% recurrence in both cohorts, spanning 11 families. Comparison is at the pathway-family level because the two cohorts frequently engage the same family through different ligands; cell-type labels were not harmonised between cohorts. Direction is not interpretable for homophilic pairs (PTPRM, JAM, CD99), where the same molecule serves as ligand and receptor and CellChat assigns both orientations.

This transcriptional phenotype was accompanied by differences in cell morphology and spatial organization. Senescent fibroblasts had larger cell and nuclear areas than regular fibroblasts in paired comparisons in both cohorts (Fig. 4e). They also co-occurred with one another more often than expected from the within-sample reference, especially at short distances. At 50 µm, the median self-co-occurrence ratio was 1.69 in breast and 4.62 in prostate metastases (Fig. 4f). A representative prostate metastasis illustrates these clusters and shows senescent fibroblasts expressing *ACTA2, COL4A1, CDKN1A*, and *SERPINE1* adjacent to cancer cells (Figs. 4g-h).

Given this spatial clustering, we asked which signaling interactions might occur within senescent fibroblast groups and between these cells and the surrounding bone niche. Predicted senescent-to-senescent interactions recurred across both cohorts, including PROS1-AXL, GAS6-AXL, PLAU-PLAUR, and TGFB3 receptor interactions (Fig. 4i). Predicted niche interactions included incoming adhesion, matrix-associated, BMP, and TGF-β signals, as well as outgoing CSF and PROS signaling toward macrophages (Fig. 4j). Together, the findings identify a spatially clustered fibroblast state with a shared secreted-gene program and nominate candidate routes for communication within these clusters and with neighboring cells.

### Expanded osteolineage with senescent phenotypes

Osteoclast abundance differed across cohorts (Kruskal–Wallis 3.5 10), with low fractions (0.5%) in NBM and substantial variation across breast and prostate bone metastases (Fig. 5a). We noticed that in cancer cohorts many maturing osteoclasts were multinucleated and therefore incorrectly split into several cells. Nucleus-guided resegmentation consolidated neighboring masks into single putative multinucleated osteoclasts, including examples with up to 18 nuclei (Fig. 5b). Counting each resegmented unit once, these units made up a larger share of the osteoclast population in the metastatic cohorts, most prominently in PCa (Fig. 5c). Thus, the cohorts differed not only in osteoclast abundance but also in the proportion of osteoclasts recovered as multinucleated units.

**Figure 5.**
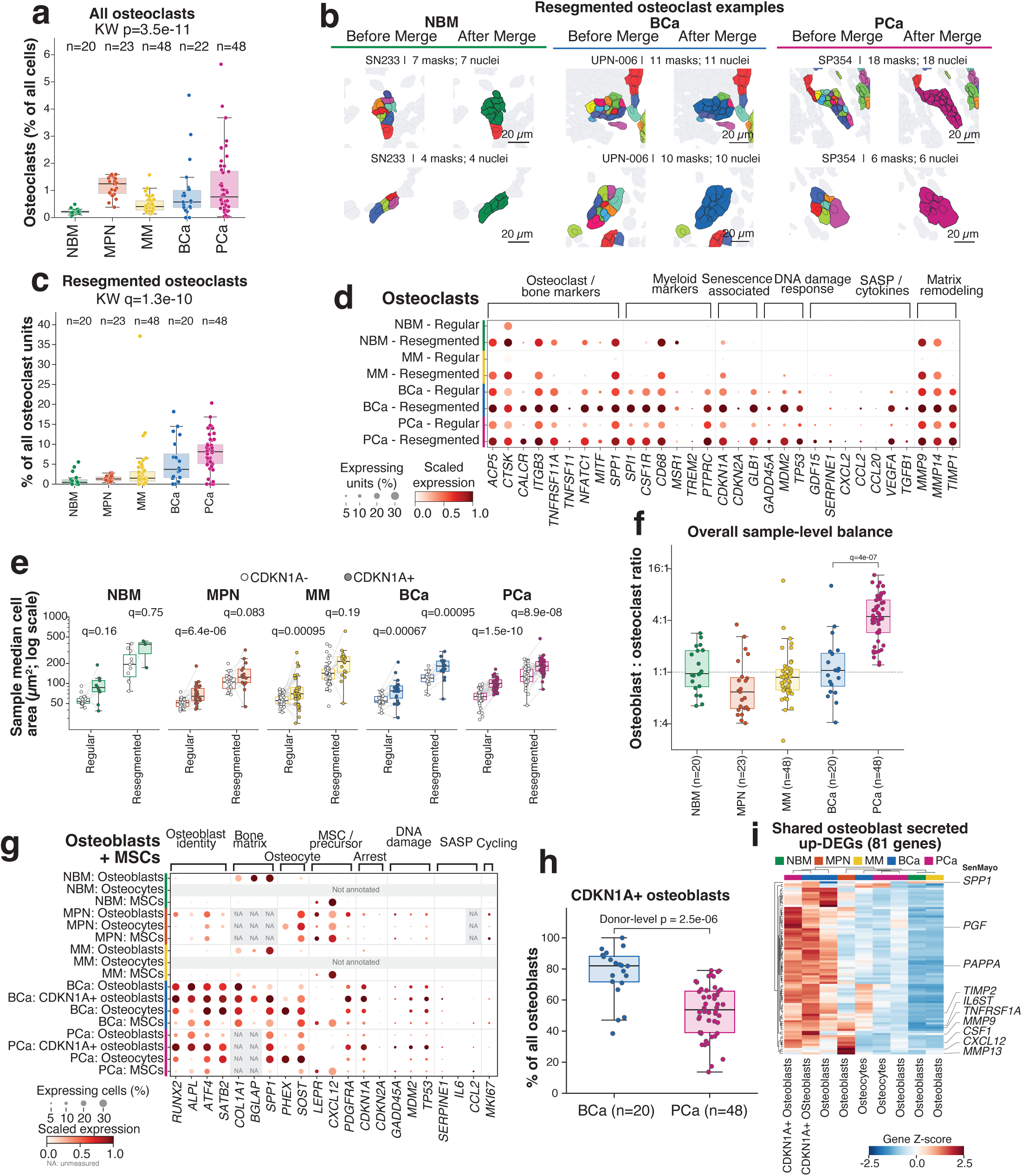
Expanded osteolineage with senescent phenotypes. a) Box plot showing percentage of all osteoclasts out of all cells in each cohort. Cancer cohorts have significantly more osteoclasts. b) Examples of resegmented osteoclasts in NBM, BCa and PCa cohorts: in left column cells are shown before resegmentation, in right columns cells are shown after resegmentation. c) Box plot showing percentage of resegmented osteoclasts out of all osteoclasts in each cohort. d) Dot plot showing expression of osteoclast and myeloid cell marker genes, cell cycle arrest genes, DNA damage response genes and SASP gene expression in resegmented and regular osteoclasts in each cohort. Resegmented osteoclasts demonstrate elevated senescent signature in BCa and PCa. e) CDKN1A+ osteoclasts tend to be larger in size in both resegmented and regular pools. f) Box plot showing osteoblast-to-osteoclast ratio across cohorts with PCa demonstrating the highest ratio among all cancer types. g) Dot plot showing expression of osteoblast, osteocyte and MSCs marker genes, cell cycle arrest genes, DNA damage response genes and SASP gene expression in osteoblasts, osteocytes, MSCs in each cohort. “NA” denotes an unmeasured gene, whereas “Not annotated” denotes a cell population not assigned in that cohort. h) Percentage of osteoblasts assigned to the CDKN1A-positive state in BCa and PCa. This is a cell-state annotation, not the individual-cell expression cutoff used in e. Points represent samples; the two-sided Mann-Whitney test was performed at the donor level 2.5 10). The denominator excludes osteocytes. i) Clustered heatmap of 81 genes encoding UniProt-annotated secreted proteins that were upregulated in the CDKN1A-positive osteoblast state in both BCa and PCa relative to the other osteoblast-lineage states in the respective cohort. Colors show gene-wise z-scores of mean expression across the displayed groups. Callouts identify genes in the SenMayo signature.

Multinucleated osteoclasts retained canonical osteoclast markers, including *ACP5*, *CTSK*, and *ITGB3*. In BCa and PCa, they also showed expression of senescence-associated, DNA damage response, and matrix-remodeling genes, including *CDKN1A*, *MDM2*, and *MMP9* (Fig. 5d). The transcriptional differences were accompanied by a morphological shift: CDKN1A-high osteoclasts had larger sample-median cell areas than CDKN1A-low osteoclasts in both the regular and resegmented categories in BCa and PCa (all 0.001; Fig. 5e). Because the size difference was also present among regular osteoclasts, it was not confined to units enlarged by merging masks. This suggests elevated *CDKN1A* expression leads to an enlarged osteoclast phenotype regardless of their maturation status.

The osteoblast compartment also showed signs of senescence. PCa metastases had a higher osteoblast-to-osteoclast ratio than BCa metastases (4 10; Fig. 5f), confirming the predominantly osteoblastic phenotype of PCa in bone. Both metastatic cancer cohorts showed the presence of *CDKN1A*-positive mature osteoblast state (Fig. 5g), which accounted for a larger fraction of osteoblasts in BCa than in PCa (donor medians, 82% versus 55%; 2.5 10; Fig. 5h). Therefore BCa mets had less overall osteoblasts than PCa but a much higher proportion of them with signs of senescence, possibly explaining the osteolytic nature of BCa mets. *CDKN1A*-positive populations retained osteoblast-lineage markers and shared 81 upregulated genes encoding annotated secreted proteins, including SenMayo genes such as *SPP1*, *CSF1*, and *MMP13* (Fig. 5g,i). The two metastatic cohorts therefore differed in cellular balance while sharing an osteoblast expression program associated with extracellular signaling and matrix remodeling.

### Metastatic cancer alters composition of nearby marrow

Next, we asked how nearby marrow cells can be affected by the presence of cancer. Marrow-rich regions accounted for most cells in NBM sections but for few or no cells in most BCa and PCa bone metastasis sections (Fig. 6a). Some metastatic specimens nevertheless retained substantial hematopoietic regions. The examples shown contained 15,653 cells in BCa and 102,061 cells in PCa, of which 82% and 84%, respectively, were hematopoietic (Fig. 6b). We asked whether marrow that persists within metastatic specimens resembles normal marrow in cellular composition and gene expression.

**Figure 6.**
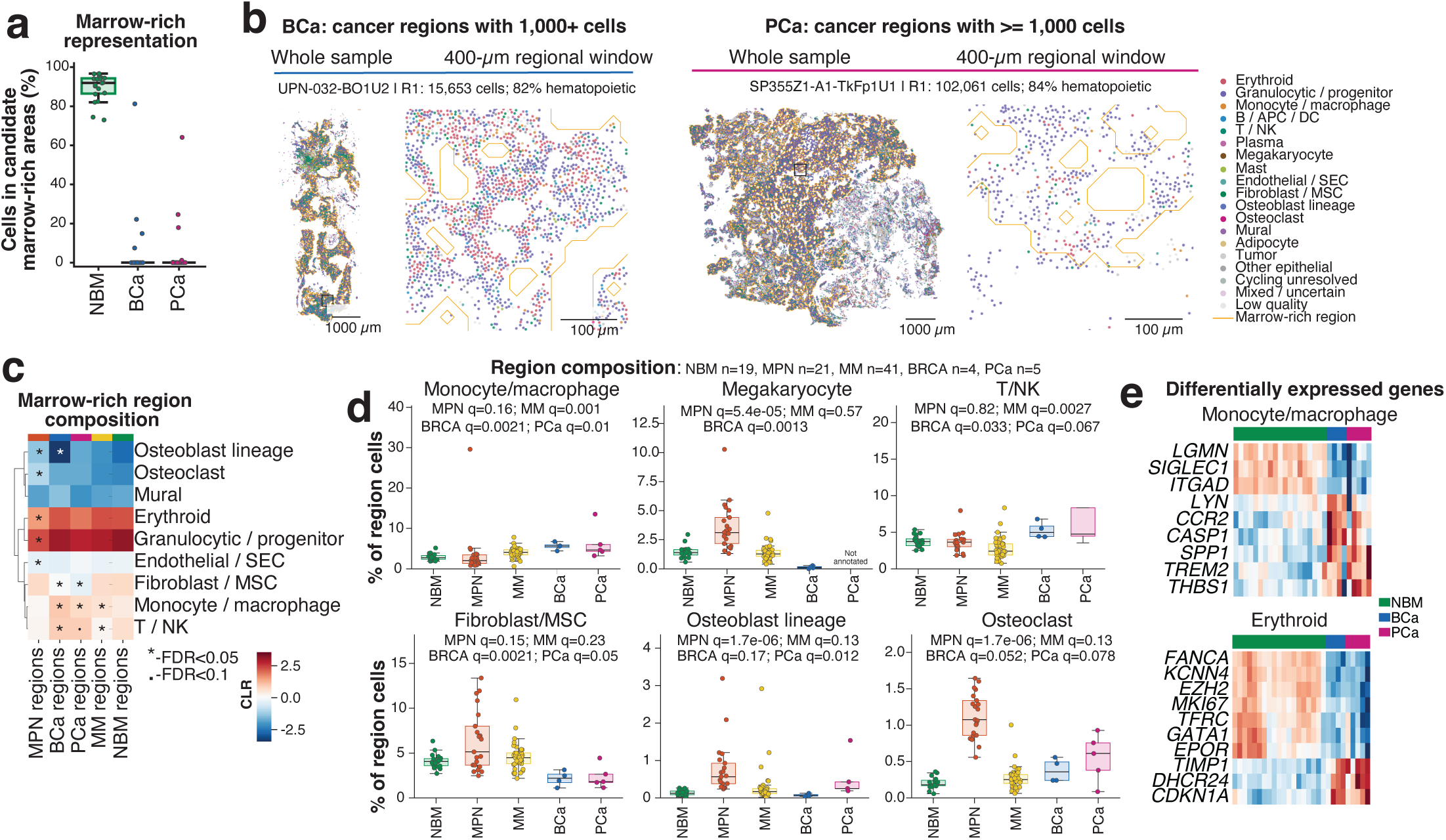
Metastatic cancer alters composition of nearby marrow. a) Percentage of all cells located within candidate marrow-rich regions in normal bone marrow (NBM), breast cancer bone metastases (BCa), and prostate cancer bone metastases (PCa). Each point represents a donor; for donors with multiple sections, section-level percentages were averaged. b) Spatial maps of representative BCa and PCa specimens containing marrow-rich regions with at least 1,000 cells. Each specimen is shown as a whole section and as a 400-µm regional window. Dots represent individual cells colored by cell type, and orange outlines delineate marrow-rich regions. The highlighted BCa and PCa regions contained 15,653 and 102,061 cells, respectively, of which 82% and 84% were hematopoietic. Scale bars, 1,000 µm for whole sections and 100 µm for regional views. c) Heatmap of marrow-rich region composition across NBM, myeloproliferative neoplasms (MPN), multiple myeloma (MM), BCa, and PCa. Colors show centered log-ratio (CLR)-transformed cohort mean cell-type proportions. Symbols indicate FDR-adjusted comparisons with NBM based on donor-level cell-type proportions: 0.05; dot, 0.05 0.10. d) Donor-level percentages of monocytes/macrophages, megakaryocytes, T/NK cells, fibroblasts/MSCs, osteoblast-lineage cells, and osteoclasts among cells in qualifying marrow-rich regions. Donor counts were NBM, 19; MPN, 21; MM, 41; BCa, 4; and PCa, 5. Each disease cohort was compared with NBM using a two-sided MannWhitney test, with Benjamini–Hochberg-adjusted values shown. Megakaryocytes were not annotated in PCa; the missing comparison does not indicate biological absence. e) Heatmaps of selected differentially expressed genes in monocytes/macrophages and erythroid cells from marrow-rich regions containing at least 1,000 cells. Differential expression was tested separately within each lineage using donor-level pseudobulk DESeq2, comparing BCa or PCa with NBM. Displayed genes met FDR 0.05 and an absolute shrunken fold change of at least 0.5 in at least one cancer-versus-NBM contrast. Columns represent individual donors, ordered NBM, BCa, and PCa; colors show gene-wise standardized expression across donors.

Despite remaining hematopoietic-rich, these regions differed from NBM in their cellular composition (Fig. 6c,d). Monocytes and macrophages accounted for a larger fraction of region cells in BCa and PCa than in NBM (donor medians: 5.6% and 4.7% versus 2.7%), whereas fibroblasts and mesenchymal stromal cells accounted for a smaller fraction (2.2% and 1.8% versus 4.1%). The pattern was not uniform across diseases: multiple myeloma also showed a higher monocyte/macrophage fraction, while MPN showed higher fractions of megakaryocytes and both osteoblast-lineage cells and osteoclasts.

The differences extended beyond cell proportions. Donor-level pseudobulk analysis showed lower expression of erythroid differentiation and proliferation genes, including *GATA1*, *EPOR*, *TFRC*, and *MKI67*, alongside higher *CDKN1A* and *TIMP1* expression in erythroid cells from both metastatic cohorts (Fig. 6e). Monocytes and macrophages likewise showed lower *SIGLEC1* and higher *SPP1* and *THBS1* in both cohorts, with significant *TREM2* upregulation in PCa, which is reported to drive osteoclastogenesis^20^. These myeloid expression changes are consistent with an altered activation or matrix-remodeling state. Together, the findings show that marrow-rich regions can persist in metastatic specimens while differing from NBM in both the proportions and transcriptional profiles of their resident cells.

## Discussion

Our findings support a model in which bone marrow aging involves coordinated changes across hematopoietic and stromal compartments, which become amplified in the context of breast and prostate cancer. In our mouse model, senescence induction recapitulated several features of physiological marrow aging, however, the limited overlap in significant age-and doxorubicin-associated genes highlights important differences between these processes. Cell-type-specific genetic models and complementary senescence-induction strategies will be needed to distinguish senescence-dependent effects from other treatment-related responses. The partial reversal effects of navitoclax may also reflect incomplete senescent-cell clearance or differential cell-type.

In human marrow, we observed inflammatory transcriptional changes in HSPCs, myeloid-skewed hematopoiesis, and pronounced remodeling of naïve CD4 T cells, alongside coordinated SASP-associated expression across stromal populations. These findings suggest that aging-associated stromal states are shaped by shared microenvironmental influences and warrant further protein-level and functional validation. The cross-sectional design and modest cohort size also limited our ability to distinguish age effects from interindividual variation, particularly for HSPC changes that showed trends but did not reach statistical significance. Larger longitudinal cohorts will be needed to resolve the temporal and functional relationships among these changes.

In breast and prostate cancer bone metastases, fibroblasts with senescence-associated expression combined activated fibroblast markers with larger cell and nuclear areas and a shared program of genes encoding secreted proteins. These fibroblasts preferentially co-occurred with one another, suggesting that the state is concentrated in local stromal patches rather than evenly distributed throughout metastatic tissue. Spatial ligand–receptor inference nominated TGF-β signaling and pericellular proteolysis as candidate mechanisms through which these patches might sustain fibrosis and influence nearby cells. The data do not establish whether these fibroblasts arise from resident fibroblasts or mesenchymal stromal cells, or whether their secreted products drive fibrosis rather than reflect an established fibrotic niche.

Osteolineage remodeling revealed a distinction between the balance of cell types and the states of cells within each lineage. Prostate metastases had a higher osteoblast-to-osteoclast ratio than breast metastases, yet the annotated CDKN1A-positive state accounted for a larger fraction of osteoblasts in breast metastases. Despite this difference in prevalence, CDKN1A-positive osteoblasts shared a secreted-gene expression program across the two cancers. In both cancers, CDKN1A-high osteoclasts were larger in the regular as well as the resegmented category, indicating that the size association was not confined to merged masks. Neither the cell-count ratio nor the expression and size differences establish rates of bone formation or resorption; those will require direct functional measurements.

The inclusion of marrow-intrinsic malignancies allowed us to test how specific these senescence-associated stromal programs are to metastatic bone disease. While MPN fibroblasts and MSCs upregulated a subset of the senescence signatures in their BCa/PCa counterparts, neither MPN nor multiple myeloma strongly expressed the secretory and matrix-remodeling programs that characterizing senescent populations in breast and prostate metastases. Osteoclasts in MPN are potentially part of the malignant clone, providing an alternative explanation why we did not observe strong senescence signatures and increase in cell size in their multinucleated osteoclast units. These absences are informative in different ways. Myelofibrosis is defined by stromal expansion and matrix deposition, yet MPN fibroblasts and mesenchymal stromal cells did not adopt the senescence-associated secretory state, indicating that marrow fibrosis can arise through mechanisms distinct from those operating in metastatic bone. Multiple myeloma is well documented to harbor inflammatory mesenchymal stromal cells^21^, yet this inflammatory activation was not accompanied by the secreted-gene and matrix program described here, suggesting that stromal inflammation and senescence-associated secretion are separable states rather than points along a single continuum. Together, these comparisons argue that the senescent fibroblast program is not a generic consequence of marrow tumor burden or fibrosis, but is instead associated with specific tumor engagement of the bone–stromal interface. Because the malignant cohorts were profiled spatially without matched functional assays, we cannot exclude comparable states existing below detection or arising at disease stages not sampled here.

Finally, the marrow-rich regions that persisted within metastatic specimens were not equivalent to normal marrow. They contained higher proportions of monocytes and macrophages and lower proportions of fibroblasts and mesenchymal stromal cells. Within these regions, erythroid cells showed reduced expression of differentiation and proliferation genes alongside higher *CDKN1A*, while monocytes and macrophages showed higher *SPP1* and *THBS1*, consistent with altered myeloid activation or matrix remodeling rather than evidence of osteoclast lineage commitment. Because relatively few metastatic donors retained large marrow-rich regions, these findings describe the marrow that remained, not all metastatic specimens. Taken together, the results point to fibrosis, altered bone-cell balance, and changes in residual hematopoietic niches as related but distinct features of metastatic marrow. A key next step is to test whether senescence-associated stromal programs help sustain these changes and whether they can be targeted without impairing normal marrow function and bone repair.

## Methods

### Study ethics and sample collection

Studies involving human participants were performed per ICH guidelines. All patient samples were obtained with informed, written consent overseen by Washington University’s Institutional Review Board. Bone marrow liquid and solid biopsies and femoral tissues were collected from healthy donors as part of the Cellular Senescence Network (SenNet) Program supported by the NIH Common Fund were collected under the IRB protocol 202108156. Additional healthy donor bone marrow aspirates and solid biopsies were collected under an institutional banking trial for health donors, IRB protocol 201103258.

### Single-cell RNA-seq sample preparation

Viably frozen aspirate samples were thawed in a PBS-based buffer (2 mM EDTA and 0.5% (w/v) bovine serum albumin; prepared by supplementing 1 L of PBS with 4.4 mL of sterile 0.5 M EDTA and 110 mL of 5% (w/v) BSA, 40% (v/v) heat-inactivated fetal calf serum, and 4.5 g/L (25 mM) glucose) and enriched for viable cells using the Miltenyi Dead Cell Removal kit per manufacturer’s instructions prior to loading onto 10X Genomics GEM platform.

Liquid slurries freshly collected from orthopedic surgery were cleared of large clots, and 20–30 mL of slurry was diluted with 20 mL ice-cold 1x PBS. All centrifugation steps were 450 rcf for 5 min at 4 °C. After an initial spin, the supernatant and floating lipid layer were aspirated and the pellet digested in 1.6 mg/mL Collagenase Type 2 for 30 min at 37 °C with shaking at 150 rpm. The digest was filtered through a 70 µm strainer and pelleted, and erythrocytes were lysed in RBC lysis buffer for 5 min at room temperature. Lysis was quenched with two volumes of PBS and the cells pelleted; this step was repeated until erythrocytes were fully removed. Cells were resuspended in 10 mL PBS, counted on a Countess automated counter, and resuspended in wash buffer (1x PBS, 10% BSA, SUPERase-In RNase inhibitor). From here, samples were either loaded onto 10X Genomics GEM platform or subdivided for sorting.

CD34⁺ cells were isolated using the EasySep Human CD34 Positive Selection Kit II (STEMCELL Technologies, #17856) according to the manufacturer’s instructions, with cells resuspended at 1×10 cells/mL in PBS with 2% fetal bovine serum and 1 mM EDTA, incubated with selection cocktail (100 µL/mL) for 10 min at room temperature, and then with RapidSpheres (75 µL/mL) for a further 5 min. Labeled suspensions were brought to 1 mL with running buffer and subjected to five rounds of magnetic separation (3 min each at room temperature), after which the enriched CD34⁺ fraction was resuspended in wash buffer at 1,500–2,000 cells/µL for downstream single-cell RNA sequencing at 1:1 ratio with unsorted fraction. For the case of CD34+/CD45-enrichment, half of the unsorted fraction of the sample was resuspended in MACS buffer with Miltenyi CD45 microbeads (20 µL per 1×10 cells), and incubated for 15 min at RT. Cells were filtered through a 70 µm strainer into sorting buffer and depleted on an AutoMACS Pro Separator (Miltenyi Biotec) using the “depletes” program; final loading for this sample was 1:1:1 unsorted:CD45-:CD34+. Sorted fractions were counted under microscopy using a hemocytometer.

Single-cell libraries were prepared using 10x Genomics Chromium Next GEM Single Cell 3′ v3 and GEM-X 3’ v4 chemistries. Per 10X Genomics guidelines, samples were loaded to target 10,000 captured cells for 3’ v3 and 20,000 cells for GEM-X 3’ v4 cDNA preparation. Full-length 3′ cDNA was enzymatically fragmented and size-selected using SPRIselect beads for library construction (10x Genomics 3′ v3 Dual Index kit; 10X Genomics Library Construction Kit C). Library quality and concentration were assessed using an Agilent TapeStation, and libraries were sequenced on NovaSeq 6000 S4 flow cells (XP workflow; 50×10×16×150 bp recipe) to a target depth of 25,000–50,000 reads per cell.

### scRNAseq data processing and QC

scRNAseq demultiplexing, alignment to human reference genome (GRCh38), and generation of gene-by-cell expression matrices were performed using Cell Ranger v6.0.1 (for Next GEM 3’ v3 chemistry) or Cell Ranger v8.0.1 (for GEM-X 3’ v4 chemistry) (10x Genomics). Per-sample CellRanger outputs were corrected for ambient RNA with SoupX, and the resulting fractional counts rounded to integers before being merged across samples (inner join on shared genes) into a single AnnData object. Doublets were called per sample with Scrublet, and cells were retained if they had ≤20% mitochondrial counts, 1,000–15,000 total counts, 200–10,000 detected genes, and were not predicted doublets. Genes detected in fewer than 3 cells were removed. Counts were normalized to median library size (excluding highly expressed genes), log1p-transformed, and the top 3,000 highly variable genes selected with sample_ID as batch key, with immunoglobulin locus genes (HGNC) excluded from HVG selection to prevent clonal plasma-cell transcripts from dominating the embedding. PCA, nearest-neighbor graph construction, UMAP, and Leiden clustering (resolution 0.5) were performed in Scanpy, and clusters were assigned to major lineages (erythroid, myeloid, T/NK, B, plasma cell, pDC, HSPC, MSC/stromal, endothelial/stromal) by marker gene expression. Cells annotated as low quality (mixed markers, low complexity) were excluded.

### scRNAseq cell type annotation

Each major lineage was subset and reprocessed independently: 2,000 highly variable genes were selected per lineage with immunoglobulin genes excluded and sample_ID as batch key, followed by PCA, Harmony integration over sample_ID, nearest-neighbor graph construction and UMAP on the Harmony-corrected embedding, and Leiden clustering across a resolution sweep (0.1–1.0). Subclusters were annotated by canonical marker expression together with Wilcoxon rank-sum differential expression restricted to protein-coding genes after removing immunoglobulin, hemoglobin, ribosomal (RPL/RPS), and mitochondrial transcripts, yielding paired broad (ct_broad) and fine (ct_fine) labels; lineages with residual internal structure, such as the neutrophil compartment, were subset and re-embedded a second time under the same procedure. Because progenitors are distributed across lineage-committed clusters, cells labeled HSPC, CLP, and MEP within the myeloid, pDC, and erythroid objects were pooled with the primary HSPC lineage, stripped of inherited embeddings and cluster assignments, and re-processed from HVG selection through Leiden clustering at higher resolution to resolve progenitor subsets and the eosinophil/basophil/mast lineage. Final annotations were propagated back to the merged object by barcode, ct_fine labels collapsed to ct_broad via a fixed mapping, and clusters failing annotation removed to produce the final object.

### Bone tissue sectioning for Xenium

Selected bone tissues were decalcified and embedded in formalin-fixed paraffin-embedded (FFPE) blocks. FFPE blocks that passed quality screening were incubated on ice for 10 to 30 minutes and sectioned at 5 μm thickness. Sections were floated on a 37 °C water bath, and then mounted onto Xenium slides. Following overnight drying at room temperature, slides were subsequently incubated at 42 °C for 3 hours before storage in a desiccator at room temperature for up to 4 weeks.

### Xenium data generation

Formalin-fixed paraffin-embedded (FFPE) blocks of interest were identified and screened by hematoxylin and eosin (H&E) staining. Selected blocks were sectioned at 5 µm and mounted on Xenium slides following the FFPE Tissue Preparation Guide (10x Genomics, CG000578; Rev C for the 477-plex assay, Rev D for Xenium Prime 5K). Slides were deparaffinized with sequential xylene and ethanol washes, then decrosslinked using the FFPE Tissue Enhancer (10x Genomics, CG000580, Rev D) for the 477-plex assay, or using the FFPE Tissue Enhancer with diluted Perm Enzyme B (modified from 10x Genomics, CG000580, Rev E) for Xenium Prime 5K.

For the 477-plex assay, overnight in situ probe hybridization was performed using the Human Multi-Tissue and Cancer Panel (10x Genomics, 1000626), targeting 377 genes; selected samples received an additional custom panel targeting 100 further genes. For Xenium Prime 5K, pre-designed and add-on custom priming oligos were first hybridized to target RNAs in situ, followed by RNase treatment to release the RNA strand and a polishing step, after which overnight probe hybridization was performed using the Xenium Prime 5K Human Pan Tissue & Pathways Panel (10x Genomics, 1000724), targeting 5,001 genes, together with a custom panel targeting 100 additional genes.

In both assays, probes were subsequently ligated and underwent rolling circle amplification, generating multiple copies of the gene-specific barcode for each RNA target. Cell segmentation reagents were applied during a staining workflow to label nuclei, membranes, and cytoplasmic regions for automated morphology-based segmentation. Background was quenched using an auto-fluorescence mixture, and nuclei were counterstained with DAPI to aid in sample tracking and cell boundary segmentation (10x Genomics, CG000749, Rev B for the 477-plex assay; CG000760, Rev C for Xenium Prime 5K).

Slides were loaded onto the 10x Xenium Analyzer (10x Genomics, 1000481) for imaging and initial data processing according to the manufacturer’s instructions (10x Genomics, CG000584, Rev F). Fluorescently labeled oligos bound to the amplified DNA probes, and iterative rounds of probe hybridization, imaging, and stripping generated optical signatures for each barcode, which were decoded to gene identities. Post-Xenium H&E staining was performed after the run as described (10x Genomics, CG000613, Rev B).

### NBM Xenium 477 data processing and cell type annotation

Per-sample Xenium outputs were loaded from cell_feature_matrix.h5, restricted to Gene Expression features, and concatenated across samples on shared genes; cell centroids, cell and nucleus area, and nucleus count were joined from each run’s cells.parquet and centroids stored as obsm[’spatial’]. Custom SNV/genotyping probes were removed from the expression matrix prior to QC. Cells were filtered using three complementary criteria: sample-adaptive lower bounds on total counts and detected genes, set at median − 5xMAD on the log1p scale with absolute floors of 5 counts and 3 genes; a per-sample Leiden clustering (QC-only) in which clusters whose third-quartile depth fell below the sample’s own first quartile were flagged as low-count; and a negative-control probe fraction (control probe counts / total counts) ≤ 0.1. Cells failing any criterion were removed, as were genes with fewer than 10 total counts. Counts were normalized to 10,000 per cell and log1p-transformed, probes detected in <0.1% of cells were excluded from feature selection, the 2,000 most variable remaining probes were used for PCA (30 components, on z-scored and clipped values), and a neighbor graph (k = 15) and UMAP were computed. An initial pre-integration Leiden sweep (resolutions 0.05–0.5), with clusters below 1,000 cells collapsed to “Unknown,” was evaluated by cluster-count elbow, per-resolution UMAPs, and marker dotplots; provisional coarse labels assigned at resolution 0.5 were used to seed semi-supervised integration. scVI was trained on raw counts with Sample_ID as batch (10 latent dimensions, 1 hidden layer), and scANVI was initialized from the trained scVI model using the provisional labels with “Unknown” as the unlabeled category. The resulting scANVI latent space was used for neighbor graph construction, UMAP, and a final Leiden sweep (resolutions 0.3 and 0.5), with per-cluster sample composition tabulated as a batch-mixing check. Final cell types were assigned at resolution 0.5 from canonical marker expression and Wilcoxon rank-sum differential expression; clusters containing mixed stromal and lymphoid populations (T/NK, fibroblast/osteoblast, MSC/fibroblast, adipocyte/endothelial) were subset and re-clustered on the scANVI latent space across resolutions 0.5–2.0 to resolve osteoblasts, fibroblasts, MSC-CAR cells, adipocytes, endothelial cells, and vSMC/pericytes, and these refined labels were written back to the merged object. Donor identifiers were parsed from sample IDs and joined to clinical metadata (age, race, sex, BMI).

### Bone metastasis Xenium data processing and annotation

Prostate and breast bone-metastasis sections were profiled with the Xenium Prime 5K human panel. The cohort contained 53 mPCa sections from 44 patients and 21 mBC sections from 18 patients. Cell-by-gene counts and cell centroids were read from the Xenium output with Seurat ReadXenium (transcript quality-value threshold 20). Centroids were stored as spatial coordinates. A cell was retained when it had at least 20 transcripts and at least 15 genes. Genes were restricted to those detected in every section, leaving 5,041 genes. Counts were normalized with LogNormalize (scale factor 10,000). Raw counts were retained. Variable genes were selected with the Seurat default (vst, 2,000 genes). Up to 5,000 cells per section were sampled by leverage score (SketchData). The sketched cells were scaled and reduced by principal-component analysis. Batch structure across sections was corrected with Harmony on the first 30 principal components, using the tissue section as the batch. Neighbors and a UMAP were computed from this Harmony embedding. Sketched cells were clustered with the Leiden algorithm (FindClusters, algorithm 4) at resolution 2.0. The Harmony model, UMAP, and cluster labels were projected from the sketched cells to all cells (ProjectIntegration and ProjectData).

Cluster markers were calculated on the sketched cells with a Wilcoxon rank-sum test. Markers were filtered by Benjamini–Hochberg adjusted P < 0.01 and average log2 fold change > 1. Each cluster was assigned manually from these markers. The final identities were tumor, T/NK cells, B cells, plasma cells, myeloid cells, granulocytes, neutrophils, erythroid cells, endothelial cells, pericytes/vSMC cells, fibroblasts, osteoblasts, osteocytes, osteoclasts, and proliferating cells. Clusters with low transcript counts, mixed lineage markers, or dominant probe noise were labeled LQ (low quality). Analyses used R 4.3.3, Seurat 5.5.1, and SeuratObject 5.4.0.

### Differential expression analysis

In scRNseq: age-associated differential expression was assessed using donor-level pseudobulk profiles and PyDESeq2 (v≥0.5). Raw counts were aggregated by donor, sample type, and cell type (UPN × sample_type), retaining protein-coding genes after excluding immunoglobulin, hemoglobin, ribosomal, and mitochondrial transcripts. Pseudobulk profiles with fewer than 10 cells were excluded, and cell types were tested only if they retained ≥8 samples spanning ≥3 distinct ages. Sample types represented by fewer than 3 samples were excluded. Genes with ≥1 CPM in at least 25% of samples were tested using a negative binomial model with age as a continuous predictor (per decade) and sample type as a covariate where applicable. Wald test p-values were adjusted using Benjamini–Hochberg correction within each cell type (FDR < 0.05). Analyses were performed independently at broad and fine cell-type annotation levels and separately in CD34⁺-enriched LT-dHSC, ST-aHSC, and MPP populations. Robustness of the top 20 age-associated genes per cell type was evaluated using Spearman correlation and covariate-adjusted OLS regression, with donor-clustered standard errors where appropriate. Outliers were identified using within-sample-type robust z-scores (|z| > 3.5), and associations were considered robust if significance (p < 0.05) and direction were maintained after outlier removal. Pathway enrichment was assessed by preranked GSEA (gseapy) using Wald statistics, MSigDB Hallmark and relevant C2.CGP gene sets (v2024.1.Hs; 15–500 genes; 2,000 permutations; FDR < 0.25).

In Xenium: given the restricted, targeted nature of Xenium gene panels, age-associated expression was assessed at the individual-gene level. For each cell type, expression was summarized as donor-level mean log1p counts per 100 cells using full-panel library size, alongside the fraction of cells detecting each gene. Donor–cell type combinations with fewer than 20 cells and cell types represented in fewer than 6 donors were excluded. Age associations were evaluated across donors using Spearman correlation, with 95% confidence intervals calculated using Fisher z transformation and the Bonett–Wright standard error. OLS slopes were calculated per decade of age, and p-values were adjusted using Benjamini– Hochberg correction across tested cell types.

### Radial neighborhood discovery

Spatial neighborhoods were defined using Xenium cell centroids and a Euclidean radius-based neighbor graph (100 µm) constructed independently for each sample. For each cell, neighboring cell-type counts were converted to within-neighborhood fractions and z-scored; cells with fewer than 30 neighbors were excluded. K-means clustering was performed across k = 2–15, with k = 6 selected using a composite criterion balancing neighborhood recurrence across samples (mean Shannon entropy of Sample_ID) and cluster distinctness (mean pairwise Euclidean distance between centroids). The resulting neighborhoods (RN1–RN6) were ranked by abundance. Cell-type enrichment was assessed using one-sided Fisher’s exact and hypergeometric tests, with Benjamini–Hochberg correction across neighborhood–cell type comparisons. Neighborhood abundance and cell-type distribution across neighborhoods were quantified per sample, averaged within donors, and correlated with age using Spearman correlation, with Benjamini–Hochberg correction.

### Cell–cell communication (LIANA)

Cell–cell communication was inferred using LIANA with the CellPhoneDB method and consensus resource. Analyses were restricted to unsorted bone slurry samples and performed independently per donor using log-normalized expression. Cell groups were defined at broad annotation resolution, except T cells, which were resolved at fine annotation. Interactions required ≥10 cells per cell type and ligand/receptor detection in ≥10% of cells. Interaction scores (lr_means) represented the mean expression of the ligand in the sender and receptor in the receiver, with complexes scored by their least-expressed subunit. T cell-associated interactions were retained if detected in ≥50% of eligible donors and tested when supported by ≥8 donors. Age associations were assessed using Spearman correlation and linear slopes per decade, with Benjamini–Hochberg correction across tested interactions.

### Fibroblast reclustering and state annotation

We reclustered stromal cells originally selected as cancer-associated fibroblasts (CAFs) in breast cancer bone metastases (BCa) and fibroblasts in prostate cancer bone metastases (PCa), independently for each cohort. Starting from raw Xenium counts in the curated cross-cohort fibroblast/MSC subset, we normalized counts to 10,000 per cell and applied ‘log1p’. In each cohort, we selected 2,000 highly variable genes using the Seurat method in Scanpy, scaled expression with a maximum value of 10, and computed 30 principal components. We corrected the principal components by sample ID using Harmony and used the resulting representation to construct a 15-nearest-neighbor graph, compute UMAP (‘min_dist=0.12’), and perform Leiden clustering. We used the resolution 0.5 clusters for manual annotation based on fibroblast, mesenchymal stromal cell (MSC), and senescence-associated marker profiles.

In BCa, cluster 6 was annotated as MSCs; clusters 7 and 8 (initially called senescent fibroblasts) and clusters 0, 1, 4, and 5 (initially called CDKN1A-positive fibroblasts) were combined into the final **Senescent fibroblasts** group. In PCa, cluster 1 was annotated as senescent fibroblasts, and clusters 2 and 6 were annotated as MSCs. Remaining clusters in each cohort were annotated as fibroblasts. The figure displays the existing Harmony-based UMAP coordinates, colored by these final labels. Senescent fibroblasts is a cluster-level transcriptional annotation, not a per-cell CDKN1A threshold or functional demonstration of senescence.

### Marker expression and stromal reference

We mapped the annotated cells back to the original cohort-specific Xenium count matrices to recover measured markers absent from the intersected feature set used for reclustering. The **Other stromal cells** reference pooled annotated osteoblast states (excluding osteocytes), endothelial cells, and sinusoidal endothelial cells from the samples represented in the fibroblast analysis. This reference was weighted by its constituent cell counts and included CDKN1A-positive states; it was not a non-senescent control. Marker counts were divided by each cell’s total counts over the shared normalization gene set, multiplied by 10,000, and ‘log1p’ transformed. Additional markers outside the normalization gene set were displayed when measured in the original cohort panel.

Genes were grouped into fibroblast identity, CAF activation, MSC-associated markers, cell-cycle arrest, DNA damage response, senescence-associated secretory phenotype (SASP), and proliferation. Dot size represents the fraction of cells with nonzero counts, capped at 30% for display. Dot color represents mean log-normalized expression over all cells in the group, including zeros, min–max scaled for each gene across the four rows within each cohort (Scanpy ‘standard_scale=’var’’). Thus, color is not an absolute cross-cohort expression comparison. DCN, LUM, and COL1A1 were not assayed in the original PCa panel and are shown as missing, not zero.

### Differential expression and UniProt annotation

Within each cancer cohort, we compared each fibroblast/MSC state with the other two states pooled, using Scanpy’s tie-corrected, cell-level Wilcoxon rank-sum test on the stored log-normalized expression matrix. Benjamini–Hochberg (BH) adjustment was applied across all tested genes for each state contrast, before filtering by secretion status. A gene was counted as an upregulated secreted DEG if its BH FDR was <0.05, its Scanpy log2 fold change was >0.25, and it was detected in at least 5% of cells in the focal state, in addition to meeting the UniProt criterion below. These tests were not pseudobulk and did not model within-patient dependence.

We defined secretion status using reviewed human UniProtKB/Swiss-Prot entries from release 2026_04. A gene qualified when it mapped unambiguously to at least one protein carrying the explicit **Secreted** keyword (‘KW-0964’). We prioritized exact primary gene symbols and accepted synonyms only when they resolved to one gene. A signal peptide or membrane annotation alone was insufficient, and unmapped or ambiguous genes were not treated as known non-secreted genes. This annotation identifies genes encoding proteins annotated as secreted.

### Shared secreted-gene expression and SenMayo annotation

The heatmap includes all 148 genes that met the upregulated secreted-DEG criteria for senescent fibroblasts in **both** BCa and PCa. For each gene, we summarized the mean stored log-normalized expression across cells in each cohort-by-state group. In addition to the three fibroblast/MSC groups in each cancer, we displayed fibroblasts and MSCs from the MPN and MM reference datasets. These references contributed expression values only, not additional differential-expression contrasts. NBM was not included. We standardized each gene across the ten displayed groups using its sample standard deviation, and hierarchically clustered genes and groups by average linkage with Euclidean distance on the uncapped Z-scores. The color scale was limited to ±2.5; missing values were not displayed as zero. Callouts mark the 20 genes that also belonged to the 125-gene SenMayo set. The callouts indicate gene-set membership, not per-cell SenMayo scores.

### Cell and nuclear area

We matched the final fibroblast annotations to raw Xenium cell summaries by sample-specific raw output folder and cell barcode. We extracted cell and nuclear segmentation-mask areas in square micrometers, deduplicated repeated raw masks, and excluded nonfinite or nonpositive values separately for each metric. MSCs were excluded. For each metric, we calculated the median area separately for regular and senescent fibroblasts within each BCa patient or PCa specimen, pooling sections belonging to the same unit. Both states had to contribute at least 20 valid cells per unit. Each plotted pair represents one of 19 BCa patients or 41 PCa specimens; connecting lines link the two state medians from that unit. We used two-sided paired Wilcoxon signed-rank tests on log2 ratios of senescent to regular fibroblast medians and applied BH correction across the four cohort-by-area comparisons.

### Spatial self-co-occurrence and representative tissue views

We computed Squidpy co-occurrence separately within each sample using the updated cell-type annotations and Xenium centroids in micrometers. We counted directed pairs within cumulative radii of 10–200 µm at 10-µm increments, excluding self-pairs and pairs at coincident coordinates. For each radius, senescent-fibroblast self-co-occurrence was the fraction of neighbors of senescent fibroblasts that were also senescent fibroblasts, divided by the fraction of all directed neighbor endpoints in the same sample and radius that were senescent fibroblasts. A ratio of 1 indicates the sample-and radius-specific reference level. Larger samples were evaluated with a dual-tree radius-count implementation checked against single-split Squidpy rather than downsampling or averaging split-specific ratios. Individual curves are shown for samples with at least two senescent fibroblasts and a defined ratio (20 BCa and 48 PCa); thick curves show the median across samples at each radius. This is a descriptive spatial statistic, not a permutation-based enrichment test.

The representative PCa field shows Xenium cell centroids colored by fibroblast state, with annotated tumor cells as a pale background. The inset displays Xenium morphology, segmentation outlines, and the spatial positions of ACTA2, COL4A1, CDKN1A, and SERPINE1 transcripts in the indicated region. The field and inset illustrate spatial context and are not additional quantitative tests of proximity or expression.

### Spatially constrained ligand–receptor inference

Cell–cell communication was inferred independently for each Xenium section with CellChat v2.2.0 (R 4.3.3) run in spatial mode. For every sample, the raw Xenium bundle (cell_feature_matrix.h5, cells.csv.gz) was matched by cell barcode to the corresponding cell-type annotation table, and analysis was restricted to the intersection of cells present in all three inputs. Cells annotated as low-quality (LQ, LQ Tumor-CAF) were removed, as were all cell types represented by fewer than 10 cells in that section, since such populations are discarded downstream by filterCommunication() in any case and their inclusion destabilises the nearest-neighbour distance estimates. Counts were normalised with CellChat::normalizeData() and a spatial CellChat object was constructed with createCellChat(datatype = “spatial”) using the Xenium centroid coordinates (spatial.factors: ratio = 1, since Xenium coordinates are already in microns; tol = 5). The human CellChat database (CellChatDB.human, 3,233 curated ligand– receptor interactions) was used in full, without restriction to a signalling class. Over-expressed genes and interactions were identified with identifyOverExpressedGenes() followed by identifyOverExpressedInteractions(), retaining 812–1,197 testable ligand–receptor pairs per section.

Communication probabilities were computed with computeCommunProb() using the trimean summary of group expression (type = “triMean”, trim = 0.1), spatial distance weighting enabled (distance.use = TRUE), an interaction range of 250 µm, a contact-dependent term with a contact range of 25 µm (contact.dependent = TRUE), the spatial adjacency gate at its default k.min = 10, and population.size = FALSE. Significance was assessed by 100 label permutations with a fixed seed; interactions with permutation p < 0.05 were retained. Communication probabilities were aggregated to the pathway level (computeCommunProbPathway()) and to the network level (aggregateNet()) at the same threshold, and the full significant interaction table was exported per sample with subsetCommunication().

Two implementation details required explicit handling. First, computeCommunProb() requires the minimum scaled inter-type distance to fall in the interval [1, 2], and the package default scale.distance = 0.01 is calibrated for spot-based data; at Xenium single-cell resolution the mean nearest-neighbour distances between cell types are an order of magnitude smaller and the default aborts the run. We therefore re-implemented CellChat:::computeRegionDistance() (using FNN::get.knnx, excluding self-matches for within-type distances) to obtain the gated mean nearest-neighbour distance matrix ahead of the call, and set scale.distance per sample so that the minimum scaled distance lands at 1.5. Where computeCommunProb() nevertheless rejected the calibrated value, the shortfall factor reported in its own error message was used to rescale with 5% headroom, up to four attempts. This parameter affects only the magnitude of the reported probabilities and cannot affect significance: the permutation loop multiplies both the observed and the permuted statistic by the same spatial factor, so it cancels in the p-value. Second, the spatial adjacency gate silently removes any cell-type pair for which fewer than k.min cells of the first type have a neighbour of the second within the interaction range; such pairs receive zero probability regardless of expression. We therefore exported, for every sample, the full pairwise mean nearest-neighbour distance matrix together with the connected/disconnected status of each pair, so that absence of a predicted interaction can be distinguished from spatial disconnection.

Of 70 sections submitted, 68 completed (48 prostate, 20 breast); 2 breast sections failed and were excluded. Completed sections contained a mean of ∼48,800 (prostate; range 6,160– 279,000) and ∼51,400 (breast; range 5,360–173,000) analysed cells across 12–18 and 22–24 cell types respectively, with minimum mean nearest-neighbour distances of 23.3 µm (prostate) and 21.8 µm (breast) on average and a mean of 1.6 of 129 and 3.8 of 258 cell-type pairs spatially disconnected. A per-sample diagnostics row was written for every section, including failures.

### Senescent fibroblast-specific interactions

To isolate signalling attributable to the senescent fibroblast state rather than to fibroblasts in general, each saved CellChat object was re-read (no model was refitted) and the full probability and p-value arrays were used directly, so that a probability of exactly zero — the ligand or receptor not expressed, or the pair spatially gated — could be distinguished from a tested but non-significant pair. The unit of test was the triplet (ligand–receptor pair × partner cell type × direction), evaluated separately for signalling from the senescent fibroblasts (fib_to_partner) and to them (partner_to_fib). Fibroblast labels were excluded from the partner set, as a fibroblast partner cannot inform state specificity.

An interaction was called senescence-exclusive in a given sample when it was significant for senescent fibroblasts (probability > 0 and p < 0.05) while the conventional fibroblast population was either not expressing the pair (probability = 0) or clearly non-significant (p > 0.2). The 0.2 margin was used in place of the nominal 0.05 so that pairs falling just short of significance in the comparator are not counted as state-specific. The mirror comparison — fibroblast-significant and senescent-fibroblast-negative — was tabulated per sample as an asymmetry control. Interaction names, pathway families and signalling class (Secreted Signaling / ECM-Receptor / Cell-Cell Contact) were annotated from CellChatDB.human.

Because the per-sample permutation test is run independently in each section, no cross-sample meta-analytic p-value was computed. Instead, reproducibility was quantified as recurrence: for each (interaction × partner × direction) key, the fraction of samples within a cohort in which it met the exclusivity criterion above. Recurrence, not effect size, is the quantity carried into all cross-cohort comparisons.

### Interactions shared between the breast and prostate cohorts

Cell-type labels were deliberately not harmonised between the two cohorts, since their annotations were produced independently and forcing a common vocabulary would conflate distinct populations. Partner labels were therefore classified as present in both annotations or specific to one, and this scope was recorded alongside every comparison; conclusions about cohort specificity were restricted to partners present in both.

Cross-cohort comparison was performed at the pathway-family level (pathway_name × partner × direction), taking for each key the maximum recurrence over the ligand–receptor pairs belonging to that family. Family-level aggregation was necessary because the two cohorts frequently engage the same pathway through different family members — for example different BMP ligands — which strict pair-level matching would score as cohort-specific. A key was called shared when its recurrence reached ≥60% of samples in both cohorts, and cohort-specific when it reached ≥60% in one cohort only; the union of these sets defines the reported comparison. For shared keys, the single most recurrent ligand–receptor pair within each cohort is reported as that cohort’s representative axis, so that the underlying family members remain visible. Autocrine (senescent-to-senescent) signalling was treated as a separate analysis using the same exclusivity criterion applied to the self-pair, and was compared across cohorts at the individual ligand–receptor level, since no family-level reconciliation was needed there.

### Osteoclast resegmentation

Within each sample, two osteoclast masks were candidate fragments of the same cell if the minimum distance between their nucleus polygons was at most 1 µm. We rejected a candidate link if a non-osteoclast cell mask intersected a corridor formed by buffering the line between the closest points on the two nuclei by 1 µm. We did not impose a cell-mask distance or centroid-distance cutoff. Connected components of the accepted links defined osteoclast units: components with at least two original masks were termed “resegmented osteoclasts”, and unmerged masks were termed “regular osteoclasts”. Resegmentation is an operational definition and does not independently establish multinucleation.

For each merged unit, raw transcript counts were summed across its constituent masks, and its spatial position was calculated as the cell-mask-area-weighted mean of their centroids. Resegmented cell area was measured from the geometric union of the cell polygons, counting overlapping area only once and without filling gaps. The number of nuclei shown in the zoomed examples was obtained by summing the Xenium nucleus counts of the constituent masks. In the paired before/after views, the same field is shown with distinct colors for the original fragments and one color for the resulting unit.

### Osteoclast abundance and resegmentation frequency

For each sample, the percentage of all cells annotated as osteoclasts was calculated as 100 × (regular + resegmented osteoclast units) / (original total cells − original osteoclast masks + resulting osteoclast units). Thus, a merged unit contributes one cell to both the numerator and denominator. Other cell types were retained in the denominator, and samples with no annotated osteoclasts were included as zero. Osteoclasts without usable boundaries were retained as singletons for this analysis. The percentage of resegmented osteoclasts used a different denominator: all resulting osteoclast units in that sample. Only samples with usable resegmentation results and at least one osteoclast were eligible for that analysis; samples with zero merges were retained. Boxplot points represent samples. The all-cell percentage was compared across cohorts using a Kruskal–Wallis test. For the resegmented percentage, the Kruskal–Wallis p-value was adjusted by Benjamini–Hochberg (BH) across three endpoints: resegmented percentage, area, and nuclei count.

### Osteoclast marker expression

Marker expression was summarized separately for regular and resegmented osteoclasts within each cohort. Counts were normalized per unit as

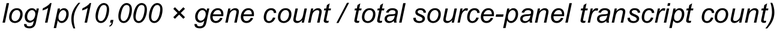

after aggregation; units with zero total counts were excluded. The dot plot used genes shared across the contributing assays. MPN was excluded from this panel because SPP1 was not available in its source gene panel. Dot size represents the percentage of units with detectable expression, with the maximum size reached at 30%. Color represents mean log-normalized expression, including zeros, min–max scaled across the displayed groups separately for each gene (Scanpy standard_scale=’var’).

### CDKN1A detection and osteoclast area

An osteoclast unit was classified as CDKN1A-positive if it contained at least one raw CDKN1A transcript, or CDKN1A-negative if none was detected. Cell area was measured from the original cell polygon for regular osteoclasts and from the polygon union for resegmented osteoclasts. Within each sample and morphology class, we calculated median cell area separately for the two expression groups. The plot displays these sample medians on a logarithmic axis and connects medians from the same sample when both groups were present. For inference, we calculated the log2 ratio of positive to negative median area within each paired sample, averaged these log2 ratios across samples in the same donor block, and tested the resulting donor-block contrasts against zero using a two-sided Wilcoxon signed-rank test. BH correction was applied across the ten cohort-by-morphology comparisons. CDKN1A detection was not used as a standalone criterion for senescence.

### Whole-section osteoblast-to-osteoclast balance

We counted osteoblasts and annotated osteoblast states, including senescent osteoblast precursors, across each sample without restricting cells to a spatial neighborhood or marrow-rich region. Osteocytes were excluded from the osteoblast numerator. The denominator included regular and resegmented osteoclasts, each counted once. In samples with at least one osteolineage cell, the plotted ratio was ‘(osteoblasts + 0.5) / (osteoclasts + 0.5)’, displayed on a log2 axis. Cohorts were compared by pairwise two-sided Mann–Whitney U tests on sample ratios with BH correction across all ten cohort pairs. This ratio measures the balance of annotated cell counts, not rates of bone formation or resorption.

### Spatial inputs and cell-type harmonization

We analyzed Xenium sections from normal bone marrow (NBM), myeloproliferative neoplasms (MPN), multiple myeloma (MM), and breast (BCa) and prostate (PCa) cancer bone metastases. Each section was analyzed separately in its Xenium coordinate system. We used the existing cohort-specific cell-type annotations, including the updated stromal states in BCa and PCa and the original coarser NBM annotations. For cross-cohort comparisons, we mapped fine annotations to broad, comparable groups. For example, B cells, antigen-presenting cells and dendritic cells were combined (B/APC/DC), fibroblasts and mesenchymal stromal cells were combined, and the osteoblast-lineage group included osteocytes. Established resegmented osteoclasts were represented by one cell unit at their merged centroid. Low-quality and ambiguous annotations were retained in total-cell denominators but did not provide evidence of hematopoietic identity. Cohort-wide absence of a cell-type annotation was distinguished from a zero count in an individual region.

### Identification of candidate marrow-rich regions

We overlaid a 25-µm square grid on each section and queried cell centroids within 100 µm of the center of every occupied tile. A tile qualified if its window contained at least 30 cell units, at least 50% of units were annotated as hematopoietic, at least five were erythroid and five were granulocytic/progenitor, and at least three hematopoietic groups were represented by at least three cells each. The hematopoietic groups included erythroid, granulocytic/progenitor, monocyte/macrophage, B/APC/DC, T/NK, plasma, megakaryocyte, and mast cells where annotated. In MPN, annotated EMP and leukemic stem cell populations formed an additional hematopoietic progenitor group.

We joined qualifying tiles using eight-neighbor connectivity, including diagonal neighbors. A connected component was retained at discovery if it contained at least four occupied tiles, 100 actual cell units, and 50 actual hematopoietic units. Cells were assigned by the tile containing their centroid, so each cell could belong to at most one region even though the 100-µm discovery windows overlap. For large-region comparisons, we applied a secondary threshold of at least 1,000 assigned cell units without changing the boundaries of retained regions. This procedure identifies candidate multilineage marrow-rich regions from annotations, not histologically confirmed normal or tumor-free marrow. The erythroid and granulocytic components are required by design, and MPN regions may include clonal hematopoiesis.

In MM, we treated all plasma-cell annotations as malignant for this analysis. These cells did not contribute to hematopoietic support or lineage-diversity criteria, but were retained in the total-cell denominator. We required a malignant plasma-cell fraction of at most 20% both in each qualifying local window and in the final connected component. Components failing the second criterion were rejected rather than trimmed. The retained regions can therefore contain residual plasma cells, and the plasma annotation alone was not used as independent evidence of malignancy.

### Section-level representation and spatial examples

For each section, we calculated the percentage of all cell units assigned to retained marrow-rich regions. The top-left plot uses the original discovery-qualified NBM regions and the ≥1,000-cell regions in BCa and PCa. For each donor, we averaged the section percentages with equal weight per section, retaining sections with no qualifying regions as zero. Each dot is one donor mean, not a pooled fraction across sections or a percentage of tissue area.

For the spatial examples, we ranked retained regions by their hematopoietic cell count and selected examples from distinct donors. The BCa section UPN-032-BO1U2 and the PCa section SP355Z1-A1-TkFp1U1 are shown as whole-section maps with candidate-region outlines and a 400 × 400 µm window around region R1. Each window was centered on the tile nearest the region’s median tile position among tiles with at least the median local cell support. Cell centroids are colored by harmonized annotation; the cell count and hematopoietic percentage above each map refer to the entire indicated region, not just the zoom window.

### Donor-level composition of large marrow-rich regions (middle row)

The five-cohort composition analysis used the same ≥1,000-cell region threshold for all cohorts, including NBM. We pooled cells across all retained regions within each section and divided each group count by the total number of assigned cell units in that section. We then averaged these percentages across the qualifying sections of each donor. Sections without qualifying regions did not contribute to conditional composition; their composition was undefined, not zero. The analysis included 19 NBM donors, 21 provisional MPN donor groups defined by CH prefixes, 41 MM donors, four BCa donors, and five PCa donors.

For each cell group, we compared donor percentages in each disease cohort with NBM using a two-sided Mann–Whitney U test, requiring at least three donors per cohort and whole-cohort annotation coverage in both groups. Benjamini–Hochberg (BH) correction was applied jointly across all 52 eligible group-by-cohort comparisons. PCa megakaryocytes were not annotated, and MM plasma cells were not compared with normal plasma cells after reclassification. The tests were unadjusted for clinical or technical covariates and are conditional on the region-selection criteria.

The composition heatmap displays nine groups with annotation coverage in all five cohorts. We averaged the section-derived percentages across donors within each cohort, added a pseudocount of 10 to the fractions, and applied the centered log-ratio (CLR) transform across the nine groups within each cohort. Both axes were hierarchically clustered on the uncapped CLR matrix (average linkage, Euclidean distance), with the displayed color scale limited to ±3.5. The figure shows only region-restricted columns; the whole-section NBM column from the source plot is not displayed. CLR colors indicate abundance relative to the geometric mean of the displayed groups within a cohort, not a fold change versus NBM or a statistical test.

### Hematopoietic pseudobulk differential expression

We restricted differential expression to hematopoietic cells inside ≥1,000-cell marrow-rich regions in BCa, PCa, and NBM. Raw integer counts for ‘Gene Expression’ features were retrieved from the original Xenium ‘cell_feature_matrix.h5’ files by exact sample-associated raw folder and cell barcode. Control features and genes not measured in every sample contributing to a given comparison were excluded. Counts were summed within each sample and broad hematopoietic group, then across samples from the same donor to create one pseudobulk per donor and group. A pseudobulk required at least 20 cells and 100 total gene-expression counts; each tested group required at least three eligible donors per cohort. Eleven comparisons qualified: six BCa versus NBM and five PCa versus NBM. BCa models included four case donors, PCa models included five except for plasma cells (four), and NBM contributed 19 reference donors. No MPN or MM cells entered these models.

We fit separate DESeq2 negative-binomial models with design ‘∼ cohort’ and NBM as the reference. Genes were required to have at least ten counts in at least three included donors. We used ‘poscounts’ size factors and two-sided Wald tests, disabled automatic count-outlier replacement, and retained DESeq2’s Cook’s-distance and independent filtering. BH-adjusted p-values were calculated within each cell-group/contrast model. Log2 fold changes were shrunk with ‘ashr’ for plotting, without replacing the Wald p-values. Volcano plots display shrunken log2 fold change versus −log10 within-model FDR. Nonflagged genes were highlighted as up or down if FDR was <0.05 and absolute shrunken log2 fold change was ≥0.5. This fold-change threshold was a display criterion, not the null hypothesis of the Wald test.

We flagged 19 tumor-associated genes for visualization using a curated epithelial/prostate marker set plus selected candidates with a median tumor-to-hematopoietic pseudobulk CPM ratio ≥8 (1-CPM pseudocount), median detection in ≥5% of tumor-labeled cells, and support from at least three donors with ≥20 tumor-labeled cells in the retained regions. Candidates qualifying in either cancer were flagged in both. Flagged genes are shown in gray and were not chosen for functional labels, but remained in the statistical models and adjusted p-value calculations. Labels were selected from curated lineage-relevant candidates passing the same FDR and effect-size cutoffs. The flag is not a spillover classifier. Differences within broad hematopoietic groups can reflect shifts in subtype composition as well as transcriptional differences within cells.

## Acknowledgements

We thank the patients, staff and scientists who contributed to this study as well as the Cellular Senescence Network (SenNet), the Human Tumor Atlas Network (HTAN) consortium, the Siteman Cancer Center, the McDonnell Genome Institute (MGI), and Departments of Surgery and Medicine at Washington University School of Medicine and the Borota gift fund and Riney Fund. We thank K. Oetjen for help with cell type annotation, J. Liu, X. Wei for assistance in data generation, E. Storrs and A. Southard-Smith for technical support.

The following grants supported this work: U54AG075934 by the National Institute on Aging to L.D., F.C., R.C.F., and S.A.S.; U2CCA233303 by the National Cancer Institute to L.D. and R.C.F.; U01CA294532 by the National Cancer Institute to L.D., F.C., and R.K.P.; U24CA211006 by the National Cancer Institute to L.D.; R01CA260112 by the National Cancer Institute to L.D. and F.C.; T32HG000045 by the National Human Genome Research Institute to A.K.

---

**Supplementary Figure 1. Marker genes used for NBM scRNA-seq cell type annotation.** a) Expression of marker genes used for broad, lineage-level cell typing.

b) Expression of marker genes used for subcluster annotation within broad cell types.

**Supplementary Figure 2. Progenitor and immune trends with aging from NBM scRNAseq.** a) Correlation of total HSPC abundance with age in unsorted samples.

b) Correlation of LT-dHSC, ST-aHSC, and MPP abundance out of total number of cells belonging to early progentor group with age in CD34+ enriched samples.

c) Top gene sets by GSEA normalized enrichment score (NES) of nominally significant age-associated DEGs in LT-dHSC, ST-aHSC, and MPP.

d) Cell type abundances correlated with age reaching nominal significance.

e) Inferred T cell interactions changing with age (LIANA).

**Supplementary Figure 3. NBM Xenium annotation and age-related trends.** a) Expression of marker genes used for Xenium 477 cell typing.

b) Correlation of vSMC/percyte and early B cell abundance with age in NBM Xenium 477 cohort.

c) Depth-adjusted % *CDKN1A*+ stromal cells per UPN.

d) Expression of marker genes used for Xenium 5K cell typing.

e) Concordance of cell type abundance in Xenium 477 versus 5K for 12 cases with both assays (left), using consensus annotations; dataset-specific cell types belonging to consensus annotations (right).

