## Supplementary Figures for "Senescence-associated stromal remodeling in bone marrow aging and hematologic and bone-metastatic diseases"

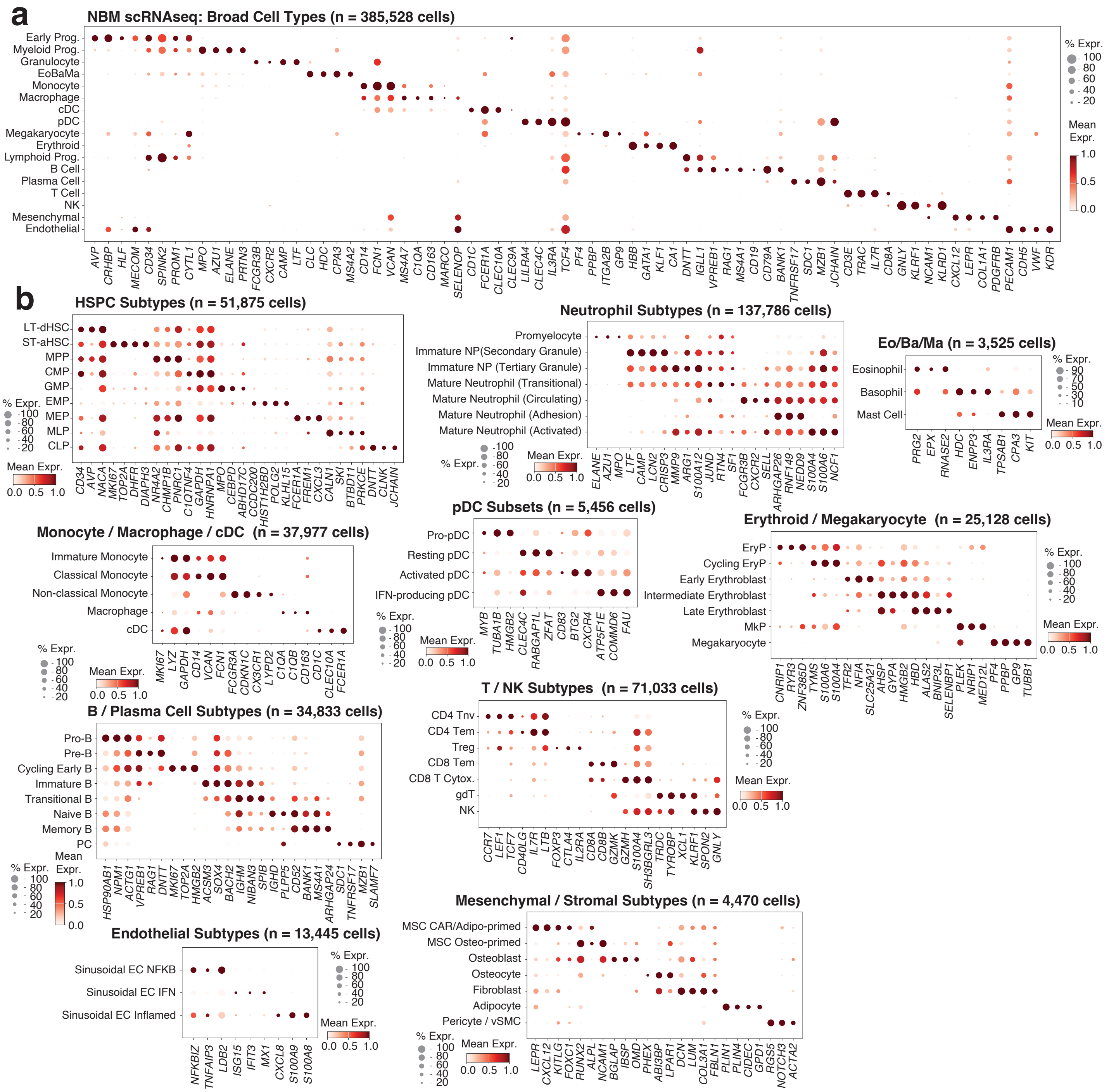
a
NBM scRNAseq: Broad Cell Types (n = 385,528 cells)
Early Prog.
Myeloid Prog.
Granulocyte
EoBaMa
Monocyte
Macrophage
cDC
pDC
Megakaryocyte
Erythroid
Lymphoid Prog.
B Cell
Plasma Cell
T Cell
NK
Mesenchymal
Endothelial
AVP CRHBP HLF MECOM CD34 SPINK2 PROM1 CYTL1 MPO AZU1 ELANE PRTN3 FCGR3B CXCR2 CAMP LTF CLC HDC CPA3 MS4A2 CD14 FCN1 VCAN MS4A7 C1QA CD163 MARCO SELENOP CD1C FCER1A CLEC10A CLEC9A LILRA4 CLEC4C IL3RA TCF4 PF4 PPBP ITGA2B GP9 HBB GATA1 KLF1 CA1 DNTT IGLL1 VPREB1 RAG1 MS4A1 CD19 CD79A BANK1 TNFRSF17 SDC1 MZB1 JCHAIN CD3E TRAC IL7R CD8A GNLY KLRF1 NCAM1 KLRD1 CXCL12 LEPR COL1A1 PDGFRB PECAM1 CDH5 VWF KDR
% Expr.
100
80
60
40
20
Mean Expr.
1.0
0.5
0.0
b
HSPC Subtypes (n = 51,875 cells)
LT-dHSC
ST-aHSC
MPP
CMP
GMP
EMP
MEP
MLP
CLP
CD34 AVP NACA MKI67 TOP2A DHFR DIAPH3 NR4A2 CHMP1B PNRC1 C1QTNF4 GAPDH HNRNPA1 MPO CEBPD ABHD17C CCDC200 HIST1H2BD POLG2 KLHL15 FCER1A FREM1 CXCL3 CALN1 SKI BTBD11 PRKCE DNTT CLNK JCHAIN
% Expr.
100
80
60
40
20
Mean Expr.
0.0 0.5 1.0
Neutrophil Subtypes (n = 137,786 cells)
Promyelocyte
Immature NP(Secondary Granule)
Immature NP (Tertiary Granule)
Mature Neutrophil (Transitional)
Mature Neutrophil (Circulating)
Mature Neutrophil (Adhesion)
Mature Neutrophil (Activated)
ELANE AZU1 MPO LTF CAMP LCN2 CRISP3 MMP9 ARG1 S100A12 JUND RTN4 SF1 FCGR3B CXCR2 SELL ARHGAP26 RNF149 NEDD9 S100A4 S100A6 NCF1
% Expr.
100
80
60
40
20
Mean Expr.
0.0 0.5 1.0
Eo/Ba/Ma (n = 3,525 cells)
Eosinophil
Basophil
Mast Cell
PRG2 EPX RNASE2 HDC ENPP3 IL3RA TPSAB1 CPA3 KIT
% Expr.
90
70
50
30
10
Mean Expr.
0.0 0.5 1.0
pDC Subsets (n = 5,456 cells)
Pro-pDC
Resting pDC
Activated pDC
IFN-producing pDC
MYB TUBA1B HMGB2 CLEC4C RABGAP1L ZFAT CD83 BTG2 CXCR4 ATP5F1E COMMD6 FAU
% Expr.
100
80
60
40
20
Mean Expr.
0.0 0.5 1.0
Monocyte / Macrophage / cDC (n = 37,977 cells)
Immature Monocyte
Classical Monocyte
Non-classical Monocyte
Macrophage
cDC
MKI67 LYZ GAPDH CD14 VCAN FCN1 FCGR3A CDKN1C CX3CR1 LYPD2 C1QA C1QB CD163 CD1C CLEC10A FCER1A
% Expr.
100
80
60
40
20
Mean Expr.
0.0 0.5 1.0
Erythroid / Megakaryocyte (n = 25,128 cells)
EryP
Cycling EryP
Early Erythroblast
Intermediate Erythroblast
Late Erythroblast
MkP
Megakaryocyte
CNRIP1 RYR3 ZNF385D TYMS S100A6 S100A4 TFR2 NFIA SLC25A21 AHSP GYPA HMGB2 HBD ALAS2 BNIP3L SELENBP1 PLEK NRIP1 MED12L PF4 PPBP GP9 TUBB1
% Expr.
100
80
60
40
20
Mean Expr.
0.0 0.5 1.0
B / Plasma Cell Subtypes (n = 34,833 cells)
Pro-B
Pre-B
Cycling Early B
Immature B
Transitional B
Naive B
Memory B
PC
HSP90AB1 NPM1 ACTG1 VPREB1 RAG1 DNTT MKI67 TOP2A HMGB2 ACSM3 SOX4 BACH2 IGHM NIBAN3 SPIB IGHD PLPP5 CD52 BANK1 MS4A1 ARHGAP24 SDC1 TNFRSF17 MZB1 SLAMF7
% Expr.
100
80
60
40
20
Mean Expr.
1.0
0.5
0.0
T / NK Subtypes (n = 71,033 cells)
CD4 Tnv
CD4 Tem
Treg
CD8 Tem
CD8 T Cytox.
gdT
NK
CCR7 LEF1 TCF7 CD40LG IL7R LTB FOXP3 CTLA4 IL2RA CD8A CD8B GZMK GZMH S100A4 SH3BGRL3 TRDC TYROBP XCL1 KLRF1 SPON2 GNLY
% Expr.
100
80
60
40
20
Mean Expr.
0.0 0.5 1.0
Endothelial Subtypes (n = 13,445 cells)
Sinusoidal EC NFKB
Sinusoidal EC IFN
Sinusoidal EC Inflamed
NFKBIZ TNFAIP3 LDB2 ISG15 IFIT3 MX1 CXCL8 S100A9 S100A8
% Expr.
100
80
60
40
20
Mean Expr.
0.0 0.5 1.0
Mesenchymal / Stromal Subtypes (n = 4,470 cells)
MSC CAR/Adipo-primed
MSC Osteo-primed
Osteoblast
Osteocyte
Fibroblast
Adipocyte
Pericyte / vSMC
LEPR CXCL12 KITLG FOXC1 RUNX2 ALPL NCAM1 BGLAP IBSP OMD PHEX ABI3BP LPAR1 DCN LUM COL3A1 FBLN1 PLIN1 PLIN4 CIDEC GPD1 RGS5 NOTCH3 ACTA2
% Expr.
100
80
60
40
20
Mean Expr.
0.0 0.5 1.0

Supplementary Figure 1. Marker genes used for NBM scRNA-seq cell type annotation.
a) Expression of marker genes used for broad, lineage-level cell typing.
b) Expression of marker genes used for subcluster annotation within broad cell types.

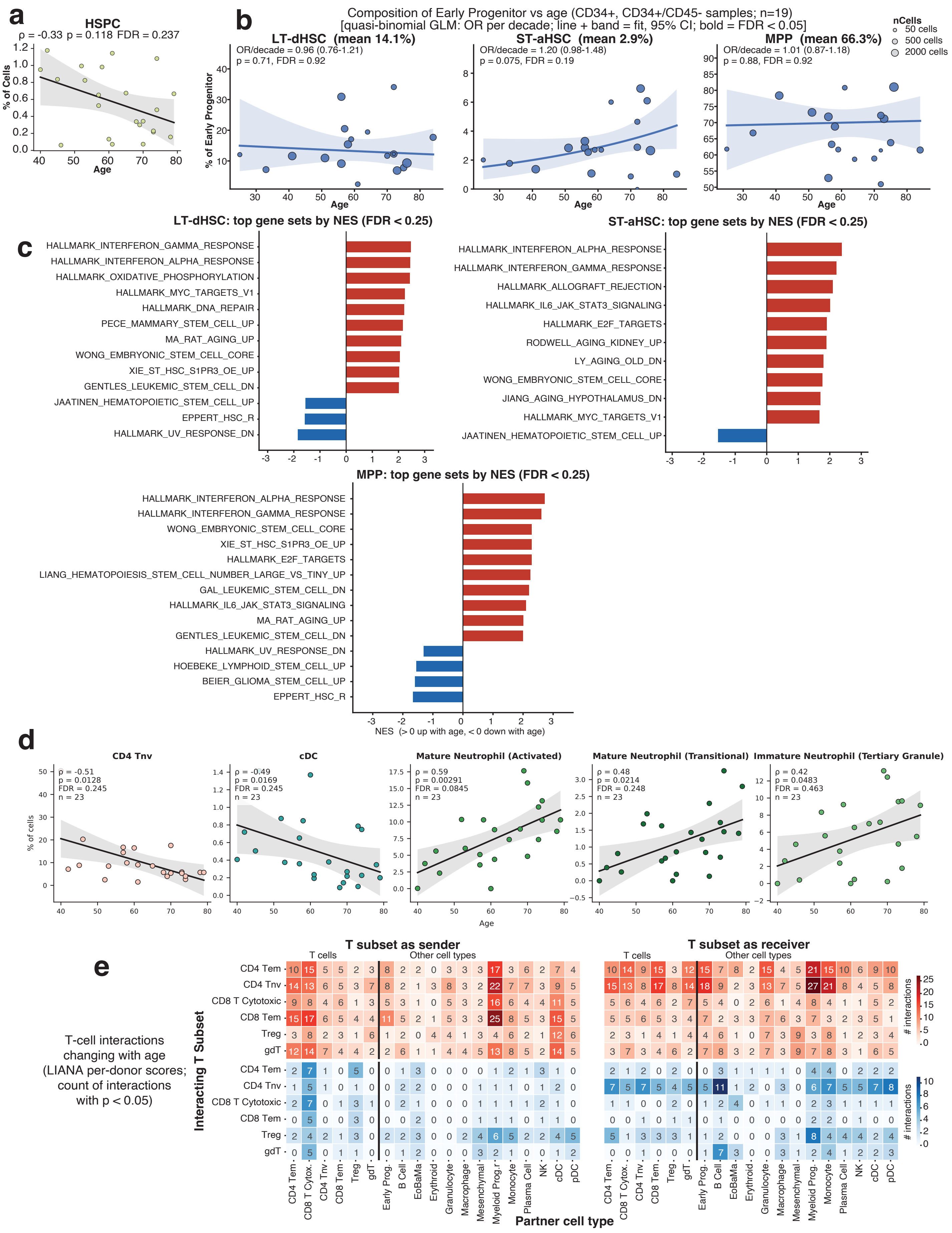
a
HSPC
ρ = -0.33 p = 0.118 FDR = 0.237
% of Cells
Age
b
Composition of Early Progenitor vs age (CD34+, CD34+/CD45- samples; n=19)
[quasi-binomial GLM: OR per decade; line + band = fit, 95% CI; bold = FDR < 0.05]
nCells
50 cells
500 cells
2000 cells
LT-dHSC (mean 14.1%)
OR/decade = 0.96 (0.76-1.21)
p = 0.71, FDR = 0.92
ST-aHSC (mean 2.9%)
OR/decade = 1.20 (0.98-1.48)
p = 0.075, FDR = 0.19
MPP (mean 66.3%)
OR/decade = 1.01 (0.87-1.18)
p = 0.88, FDR = 0.92
% of Early Progenitor
Age
c
LT-dHSC: top gene sets by NES (FDR < 0.25)
HALLMARK_INTERFERON_GAMMA_RESPONSE
HALLMARK_INTERFERON_ALPHA_RESPONSE
HALLMARK_OXIDATIVE_PHOSPHORYLATION
HALLMARK_MYC_TARGETS_V1
HALLMARK_DNA_REPAIR
PECE_MAMMARY_STEM_CELL_UP
MA_RAT_AGING_UP
WONG_EMBRYONIC_STEM_CELL_CORE
XIE_ST_HSC_S1PR3_OE_UP
GENTLES_LEUKEMIC_STEM_CELL_DN
JAATINEN_HEMATOPOIETIC_STEM_CELL_UP
EPPERT_HSC_R
HALLMARK_UV_RESPONSE_DN
ST-aHSC: top gene sets by NES (FDR < 0.25)
HALLMARK_INTERFERON_ALPHA_RESPONSE
HALLMARK_INTERFERON_GAMMA_RESPONSE
HALLMARK_ALLOGRAFT_REJECTION
HALLMARK_IL6_JAK_STAT3_SIGNALING
HALLMARK_E2F_TARGETS
RODWELL_AGING_KIDNEY_UP
LY_AGING_OLD_DN
WONG_EMBRYONIC_STEM_CELL_CORE
JIANG_AGING_HYPOTHALAMUS_DN
HALLMARK_MYC_TARGETS_V1
JAATINEN_HEMATOPOIETIC_STEM_CELL_UP
MPP: top gene sets by NES (FDR < 0.25)
HALLMARK_INTERFERON_ALPHA_RESPONSE
HALLMARK_INTERFERON_GAMMA_RESPONSE
WONG_EMBRYONIC_STEM_CELL_CORE
XIE_ST_HSC_S1PR3_OE_UP
HALLMARK_E2F_TARGETS
LIANG_HEMATOPOIESIS_STEM_CELL_NUMBER_LARGE_VS_TINY_UP
GAL_LEUKEMIC_STEM_CELL_DN
HALLMARK_IL6_JAK_STAT3_SIGNALING
MA_RAT_AGING_UP
GENTLES_LEUKEMIC_STEM_CELL_DN
HALLMARK_UV_RESPONSE_DN
HOEBEKE_LYMPHOID_STEM_CELL_UP
BEIER_GLIOMA_STEM_CELL_UP
EPPERT_HSC_R
NES (> 0 up with age, < 0 down with age)
d
CD4 Tnv
ρ = -0.51
p = 0.0128
FDR = 0.245
n = 23
cDC
ρ = -0.49
p = 0.0169
FDR = 0.245
n = 23
Mature Neutrophil (Activated)
ρ = 0.59
p = 0.00291
FDR = 0.0845
n = 23
Mature Neutrophil (Transitional)
ρ = 0.42
p = 0.0214
FDR = 0.248
n = 23
Immature Neutrophil (Tertiary Granule)
ρ = 0.42
p = 0.0483
FDR = 0.463
n = 23
% of cells
Age
e
T-cell interactions changing with age (LIANA per-donor scores; count of interactions with p < 0.05)
T subset as sender
T subset as receiver
T cells
Other cell types
Interacting T Subset
CD4 Tem
CD4 Tnv
CD8 T Cytotoxic
CD8 Tem
Treg
gdT
CD4 Tem
CD8 T Cytox.
CD4 Tnv
CD8 Tem
Treg
gdT
Early Prog.
B Cell
EoBaMa
Erythroid
Granulocyte
Macrophage
Mesenchymal
Myeloid Prog.
Monocyte
Plasma Cell
NK
cDC
pDC
Partner cell type
### interactions

Supplementary Figure 2. Progenitor and immune trends with aging from NBM scRNAseq.
a) Correlation of total HSPC abundance with age in unsorted samples.
b) Correlation of LT-dHSC, ST-aHSC, and MPP abundance out of total number of cells belonging to early progentor group with age in CD34+ enriched samples.
c) Top gene sets by GSEA normalized enrichment score (NES) of nominally significant age-associated DEGs in LT-dHSC, ST-aHSC, and MPP.
d) Cell type abundances correlated with age reaching nominal significance.
e) Inferred T cell interactions changing with age (LIANA).

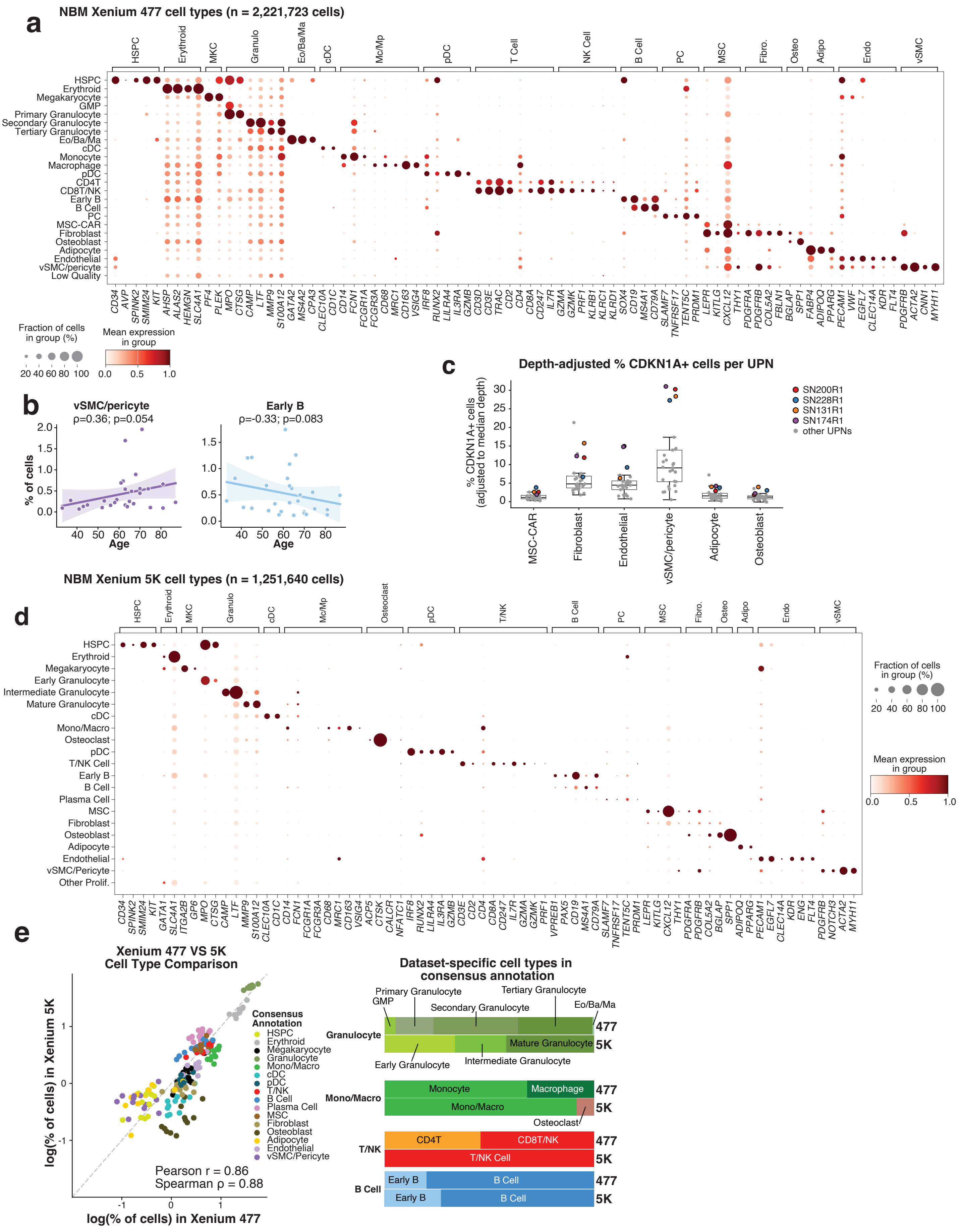
a
NBM Xenium 477 cell types (n = 2,221,723 cells)
HSPC
Erythroid
MKC
Granulo
Eo/Ba/Ma
cDC
Mc/Mp
pDC
T Cell
NK Cell
B Cell
PC
MSC
Fibro.
Osteo
Adipo
Endo
vSMC
HSPC
Erythroid
Megakaryocyte
GMP
Primary Granulocyte
Secondary Granulocyte
Tertiary Granulocyte
Eo/Ba/Ma
cDC
Monocyte
Macrophage
pDC
CD4T
CD8T/NK
Early B
B Cell
PC
MSC-CAR
Fibroblast
Osteoblast
Adipocyte
Endothelial
vSMC/pericyte
Low Quality
CD34 AVP SPINK2 SMIM24 KIT AHSP ALAS2 HEMGN SLC4A1 PF4 PLEK MPO CTSG CAMP LTF MMP9 S100A12 GATA2 MS4A2 CPA3 CLEC10A CD1C CD14 FCN1 FCGR1A FCGR3A CD68 MRC1 CD163 VSIG4 IRF8 RUNX2 LILRA4 IL3RA GZMB CD3D CD3E TRAC CD2 CD4 CD8A CD247 IL7R GZMA GZMK PRF1 KLRB1 KLRC1 KLRD1 SOX4 CD19 MS4A1 CD79A SLAMF7 TNFRSF17 TENT5C PRDM1 LEPR KITLG CXCL12 THY1 PDGFRA PDGFRB COL5A2 FBLN1 BGLAP SPP1 FABP4 ADIPOQ PPARG PECAM1 VWF EGFL7 CLEC14A KDR FLT4 PDGFRB ACTA2 CNN1 MYH11
Fraction of cells in group (%)
20 40 60 80 100
Mean expression in group
0.0 0.5 1.0
b
vSMC/pericyte
ρ=0.36; p=0.054
Early B
ρ=-0.33; p=0.083
% of cells
Age
c
Depth-adjusted % CDKN1A+ cells per UPN
% CDKN1A+ cells (adjusted to median depth)
SN200R1
SN228R1
SN131R1
SN174R1
other UPNs
MSC-CAR
Fibroblast
Endothelial
vSMC/pericyte
Adipocyte
Osteoblast
d
NBM Xenium 5K cell types (n = 1,251,640 cells)
HSPC
Erythroid
MKC
Granulo
cDC
Mc/Mp
Osteoclast
pDC
T/NK
B Cell
PC
MSC
Fibro.
Osteo
Adipo
Endo
vSMC
HSPC
Erythroid
Megakaryocyte
Early Granulocyte
Intermediate Granulocyte
Mature Granulocyte
cDC
Mono/Macro
Osteoclast
pDC
T/NK Cell
Early B
B Cell
Plasma Cell
MSC
Fibroblast
Osteoblast
Adipocyte
Endothelial
vSMC/Pericyte
Other Prolif.
CD34 SPINK2 SMIM24 KIT GATA1 SLC4A1 ITGA2B GP6 MPO CTSG CAMP LTF MMP9 S100A12 CLEC10A CD1C CD14 FCN1 FCGR1A FCGR3A CD68 MRC1 CD163 VSIG4 ACP5 CTSK CALCR NFATC1 IRF8 RUNX2 LILRA4 IL3RA GZMB CD3E CD2 CD4 CD8A CD247 IL7R GZMA GZMK PRF1 VPREB1 PAX5 CD19 MS4A1 CD79A SLAMF7 TNFRSF17 TENT5C PRDM1 LEPR KITLG CXCL12 THY1 PDGFRA PDGFRB COL5A2 BGLAP SPP1 ADIPOQ PPARG PECAM1 EGFL7 CLEC14A KDR ENG FLT4 PDGFRB NOTCH3 ACTA2 MYH11
Fraction of cells in group (%)
20 40 60 80 100
Mean expression in group
0.0 0.5 1.0
e
Xenium 477 VS 5K
Cell Type Comparison
log(% of cells) in Xenium 5K
log(% of cells) in Xenium 477
Pearson r = 0.86
Spearman ρ = 0.88
Consensus Annotation
HSPC
Erythroid
Megakaryocyte
Granulocyte
Mono/Macro
cDC
pDC
T/NK
B Cell
Plasma Cell
MSC
Fibroblast
Osteoblast
Adipocyte
Endothelial
vSMC/Pericyte
Dataset-specific cell types in consensus annotation
Granulocyte
Primary Granulocyte
GMP
Tertiary Granulocyte
Secondary Granulocyte
Eo/Ba/Ma
Mature Granulocyte
Early Granulocyte
Intermediate Granulocyte
477
5K
Mono/Macro
Monocyte
Macrophage
Mono/Macro
Osteoclast
T/NK
CD4T
CD8T/NK
T/NK Cell
B Cell
Early B
B Cell

Supplementary Figure 3. NBM Xenium annotation and age-related trends.
a) Expression of marker genes used for Xenium 477 cell typing.
b) Correlation of vSMC/percyte and early B cell abundance with age in NBM Xenium 477 cohort.
c) Depth-adjusted % CDKN1A+ stromal cells per UPN.
d) Expression of marker genes used for Xenium 5K cell typing.
e) Concordance of cell type abundance in Xenium 477 versus 5K for 12 cases with both assays (left), using consensus annotations; dataset-specific cell types belonging to consensus annotations (right).
